# A new *Plasmodium vivax* reference genome for South American isolates

**DOI:** 10.1101/2023.03.14.532329

**Authors:** Katlijn De Meulenaere, Bart Cuypers, Dionicia Gamboa, Kris Laukens, Anna Rosanas-Urgell

**Affiliations:** Department of Biomedical Sciences, Institute of Tropical Medicine Antwerp, Antwerp, Belgium; Department of Computer Science, University of Antwerp, Antwerp, Belgium; Instituto de Medicina Tropical Alexander von Humboldt, Universidad Peruana Cayetano Heredia, Lima, Peru; Departamento de Ciencias Celulares y Moleculares, Facultad de Ciencias y Filosofía, Universidad Peruana Cayetano Heredia, Lima, Peru

**Keywords:** *Plasmodium vivax*, genome assembly, PacBio sequencing, reference genome

## Abstract

**Background:** *Plasmodium vivax* is the second most important cause of human malaria worldwide, and accounts for the majority of malaria cases in South America. A high-quality reference genome exists for Papua Indonesia (PvP01) and Thailand (PvW1), but is lacking for South America. A reference genome specifically for South America would be beneficial though, as *P. vivax* is a genetically diverse parasite with geographical clustering.

**Results:** This study presents a new high-quality assembly of a South American *P. vivax* isolate, referred to as PvPAM. The genome was obtained from a low input patient sample from the Peruvian Amazon and sequenced using PacBio technology, resulting in a highly complete assembly with 6497 functional genes. Telomeric ends were present in 17 out of 28 chromosomal ends, and additional (sub)telomeric regions are present in 12 unassigned contigs. A comparison of multigene families between PvPAM and the PvP01 genome revealed remarkable variation in *vir* genes, and the presence of merozoite surface proteins (MSP) 3.6 and 3.7. Three *dhfr* and *dhps* drug resistance associated mutations are present in PvPAM, similar to those found in other Peruvian isolates. Mapping of publicly available South American whole genome sequencing (WGS) data to PvPAM resulted in significantly fewer variants and truncated reads compared to the use of PvP01 or PvW1 as reference genomes. To minimize the number of core genome variants in non-South American samples, PvW1 is most suited for Southeast Asian isolates, both PvPAM and PvW1 are suited for South Asian isolates, and PvPAM is recommended for African isolates. Interestingly, non-South American samples still contained the least subtelomeric variants when mapped to PvPAM, indicating high quality of the PvPAM subtelomeric regions.

**Conclusions:** Our findings show that the PvPAM reference genome more accurately represents South American *P. vivax* isolates in comparison to PvP01 and PvW1. In addition, PvPAM has a high level of completeness, and contains a similar number of annotated genes as PvP01 or PvW1. The PvPAM genome therefore will be a valuable resource to improve future genomic analyses on *P. vivax* isolates from the South American continent.

## Background

*Plasmodium vivax* is the second most common cause of malaria in humans after *P. falciparum. P. vivax* is widespread outside of Africa, accounting there for one-third of the malaria infections (WHO, 2021). Due to the absence of a continuous culturing system (Bermúdez et al., 2018) and the former assumption that *P. vivax* infections are benign (Howes et al., 2016), *P. vivax* research has been lagging behind that of *P. falciparum*. Only in 2008, the first *P. vivax* reference genome, Salvador I, was assembled (Carlton et al., 2008), while the *P. falciparum* 3D7 reference genome was already published in 2002 (Gardner et al., 2002).

Hitherto, several *P. vivax* genomes have been assembled for *P. vivax*. The Salvador I (PvSalI) genome was assembled from paired-read Sanger sequencing data, coming from an El Salvador isolate that was passaged in *Saimiri* monkeys (Collins et al., 1972;Carlton et al., 2008). However, it contains >2000 unassigned contigs that were not assigned to a chromosome, making it less suited for alignment of whole genome sequencing (WGS) data. Similarly, Illumina-based assemblies from four monkey-passaged isolates from Brazil, India, Mauritania and North Korea (Neafsey et al., 2012) and 1 Cambodian field isolate (Hester et al., 2013) were highly fragmented, with >1500 scaffolds. More recently, Auburn et al. (2016) released a high-quality reference genome from an Illumina-sequenced Papua Indonesian isolate (PvP01), along with two draft assemblies from Thailand (PvT01) and China (PvC01). Finally, a Thai isolate used in controlled human infections was long-read sequenced with PacBio, resulting in the high-quality PvW1 reference genome (Minassian et al., 2021).

South America is not yet represented by the existing high-quality reference genomes (PvP01, PvW1), although the majority of the malaria cases in this continent are *P. vivax* (WHO, 2021). Since *P. vivax* is genetically more diverse than *P. falciparum* (Neafsey et al., 2012;Hupalo et al., 2016;Fola et al., 2017), there is a strong geographical differentiation between isolates from different continents (Koepfli et al., 2015;Hupalo et al., 2016;Benavente et al., 2021). The PvP01 and PvW1 assemblies originate from Papua Indonesia and Thailand, and isolates from these countries are genetically distant from the South American ones (Benavente et al., 2021). Therefore, a new *P. vivax* reference genome from South American origin could improve variant calling of South American isolates and provide more accurate information on indels (insertions and deletions) and large structural variants. It could be especially beneficial to study the subtelomeric regions of South American isolates, as subtelomeres are highly dynamic with frequent duplication and recombination events (Mok et al., 2008), and contain important multigene families involved in immune evasion and host-parasite interactions (Carlton et al., 2008).

Since *de novo* assembly of a reference genome makes use of the overlaps between sequenced fragments (reads), longer reads improve confidence and quality of the assembly. In addition, long reads can better resolve repetitive or low-complexity regions often found in the subtelomeres. Short-read technologies like Illumina typically have read lengths of 50-300 bases, while Pacific Biosciences SMRT technology (PacBio) and Oxford Nanopore technology (ONT), the leading long-read sequencing technologies to date, readily exceed 10,000 bases. Therefore, PacBio and Nanopore are more suited for reference genome sequencing. At the moment of writing, PacBio sequencing has an accuracy of 99.8% (Wenger et al., 2019), similar to what can be achieved with Illumina sequencing, and higher than the 99% that can be reached with the Nanopore R10.4 chemistry (Sereika et al., 2022). Due to PacBio’s use of circular consensus sequencing (a circular read is sequenced multiple times), random sequencing errors are largely removed (Wenger et al., 2019). Although a bias for indels at homopolymer regions remains (Wenger et al., 2019;Karst et al., 2021), it is the most robust long-read sequencing method to date, which was also used to construct the Thai PvW1 reference (Minassian et al., 2021). The PvW1 sample was obtained from controlled human malaria infections, which allows extraction of a high amount of input DNA. However, low input protocols are becoming more common, and PacBio recently released their Ultra-Low DNA Input protocol (5-20 ng DNA) (PacBio, 2020b). This provides opportunities to long-read sequence *P. vivax* DNA from low blood volume and/or low parasitaemia patient samples, which is common in natural human infections.

In this study, a leukocyte-depleted blood sample from a *P. vivax* patient from the Peruvian Amazon was sequenced with PacBio using the Ultra-Low DNA Input protocol, allowing the assembly of a new and high-quality South American reference genome named PvPAM (*P. vivax* Peruvian AMazon). The PvPAM chromosomes have a high level of completeness (telomeric repeats in 17/28 chromosomal ends), although 12 unassigned contigs remain. Publicly available South American WGS data were mapped to PvPAM to provide an accurate overview of South American *P. vivax* variants. This mapping resulted in significantly less core genome variants compared to using PvP01 or PvW1 as reference genomes, indicating that PvPAM better represents South American isolates. The same was also observed for African isolates, while Southeast Asian isolates had the lowest number of core genome variants when mapped to PvW1. In the subtelomeric regions, the use of PvPAM as a reference genome resulted in a significantly lower number of variants for WGS data from all continents, outperforming PvP01 and PvW1.

## Results

### Construction of a high-quality reference genome from a low-input *P. vivax* patient DNA sample

A new reference genome was constructed from a monoclonal *P. vivax* patient sample originating from the Peruvian Amazon basin (Iquitos), which we named PvPAM (*P. vivax* Peruvian AMazon). Although the input sample contained a very low amount of parasite DNA (9 ng), the PacBio Ultra-Low Input protocol for library preparation allowed to sequence the *P. vivax* genome in high depth and enabled to build a high-quality reference genome for the South American Amazon region.

497 billion raw continuous long read (CLR) bases were sequenced, and after pre-processing, 24.5 billion high fidelity circular consensus sequencing (CCS) read bases were left, with reads being 10,670 bases long on average. When mapped to the PvP01 reference genome, this resulted in an average sequencing depth of 821x. This is well above the recommended 15x depth for assembly of a haploid genome (PacBio, 2020a), but due to the repetitive and AT-rich subtelomeric regions, higher depths are necessary to fully assemble the eukaryotic *P. vivax* genome.

Genome assembly and subsequent scaffolding resulted in 14 chromosomes, of which 4 were completely assembled, while the other 10 were scaffolds consisting of 2-4 contigs. The gaps between those contigs were patched based on the PvP01 and PvW1 reference genomes (Auburn et al., 2016;Minassian et al., 2021). The patches had a median length of 388 bases (range 41-2592 bases) (Supplementary Figure 1), which indicates that the initial PvPAM assembly was of good quality and only few bases were uncovered. Further polishing of the genome with Illumina reads, obtained in parallel from the PvPAM isolate, introduced some PvPAM-specific changes in the patched regions. In addition to the 14 chromosomes, 12 contigs could not be uniquely assigned to any chromosome, although they were identified as subtelomeric based on their low GC content (<25%) and presence of *vir* genes.

PvPAM chromosome-completeness was assessed by the presence of telomeric CCCT(A/G)AA repeats (Ponzi et al., 1985;Vernick and McCutchan, 1988), which indicate that the chromosome was fully assembled up to the 5’ or 3’ end. Telomeric repeats were only observed at one chromosomal end in PvP01, while they are present in 17/28 chromosomal ends in the PvPAM, indicating a higher number of completely assembled chromosomes (Figure 1). Eight additional telomeric repeat sequences could be found in the 12 unassigned and subtelomeric PvPAM contigs.

**Figure 1.**
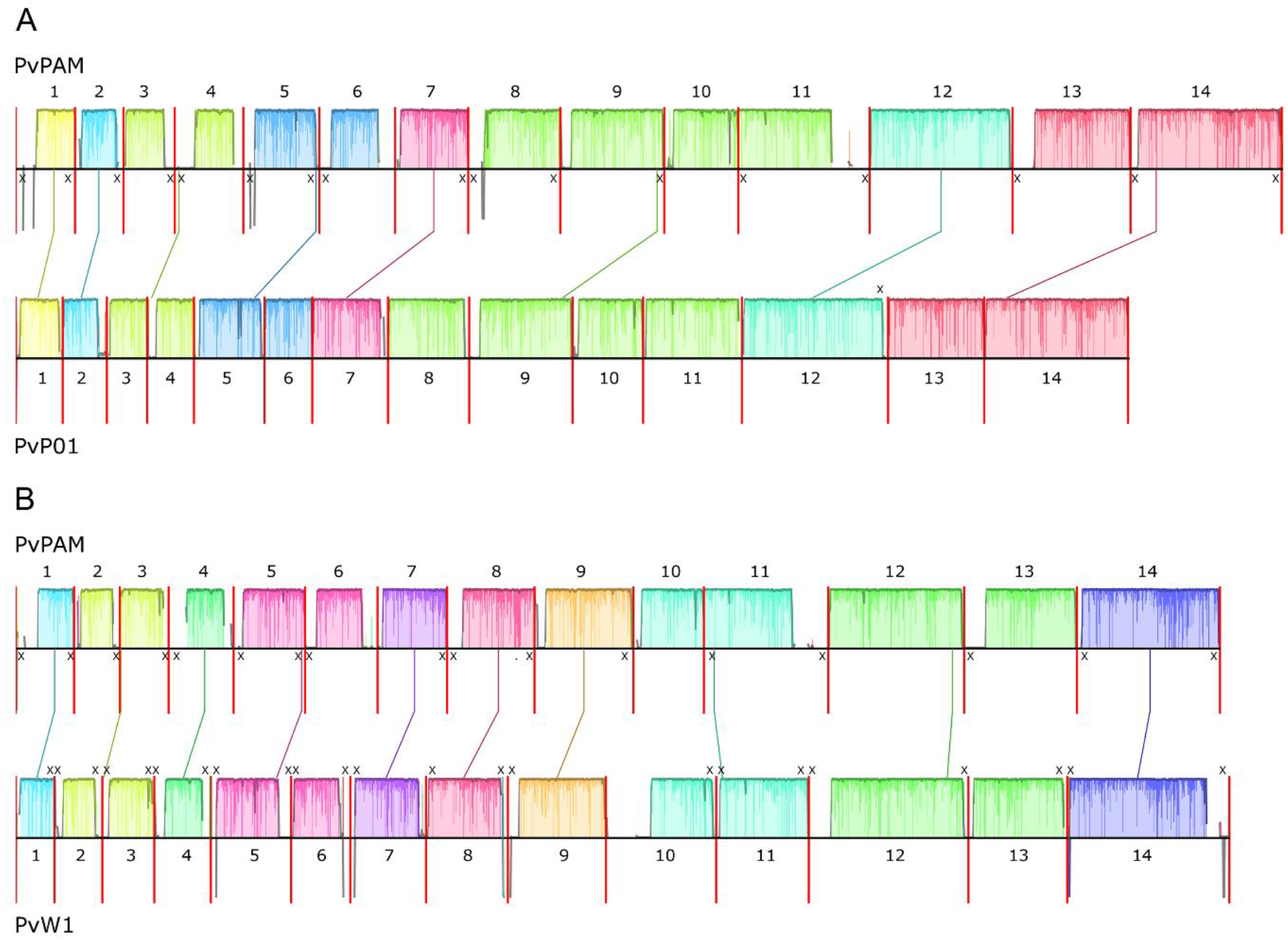
Whole genome alignments of PvPAM and PvP01 (PlasmoDB, v51) (**A**), and PvPAM and PvW1 (PlasmoDB, v60) (**B**), constructed with the Mauve Contig Mover (MCM) algorithm (Darling et al., 2010). Coloured blocks indicate homologous regions, and their height indicates the level of similarity between both reference genomes. Blank regions indicate sequences that do not align between the two reference genomes. Chromosomes are separated by a red vertical line and are numbered. If a telomeric repeat is present at the chromosomal end, this is indicated by a cross. The mitochondrial genome, apicoplast genome, and unassigned contigs are not shown.

### Structural comparison of PvPAM to the PvP01 and PvW1 reference genomes

PvPAM was compared on a structural and annotation level to the two highest quality *P. vivax* reference genomes existing to date: PvP01 (Auburn et al., 2016) and the PacBio-based PvW1 (Minassian et al., 2021). These three reference genomes have overall similar genome sizes (Table 1). However, whole genome alignments of the reference genomes reveal that most PvPAM chromosomes contain longer subtelomeric regions than their PvP01 counterparts (Figure 1A). This is because large parts of the PvP01 subtelomeric regions are divided over many short contigs that are not assigned to any chromosome, while the PvPAM reference genome is less fragmented. The subtelomeric regions of the PvW1 chromosomes are overall of similar length as in PvPAM (Table 1), but at the chromosome level, the subtelomeres often have different lengths or do not fully align (Figure 1B). We determined core genome boundaries based on GC-content and sequence identity between PvPAM, PvP01 and PvW1. These borders show that the PvW1 and PvPAM chromosomes possess longer subtelomeric regions than the PvP01 chromosomes, due to higher contiguity in PacBio-based genomes (Table 1).

**Table 1.** Comparison of the PvPAM, PvP01 (PlasmoDB, v51) and PvW1 (PlasmoDB, v60) assemblies. The assembly size includes unassigned contigs. The subtelomeric region size of chromosomes is based on the subtelomeric boundaries as defined in the Methods section. Functional genes are all genes that are not annotated as pseudogenes. PvW1 pseudogenes are not annotated and therefore unknown. Mb=megabases, kb=kilobases.

|  |  | PvPAM |  | PvP01 |  | PvW1 |  |
| --- | --- | --- | --- | --- | --- | --- | --- |
| Nuclear genome |  |  |  |  |  |  |  |
| Assembly size (Mb) |  | 29.40 |  | 29.00 |  | 28.96 |  |
| GC content (%) |  | 39.59 |  | 39.81 |  | 39.89 |  |
| Number of scaffolds assigned to a chromosome |  | 14 |  | 14 |  | 14 |  |
| Number of unassigned contigs |  | 12 |  | 226 |  | 3 |  |
| Size of unassigned contigs (Mb) |  | 1.88 |  | 4.79 |  | 1.18 |  |
| Subtelomeric genome size (Mb) | Unassigned contig | 9.41 | 1.88 | 9.01 | 4.76 | 8.98 | 1.18 |
|  | Part of chromosome |  | 7.53 |  | 4.25 |  | 7.80 |
| Number of functional genes |  | 6456 |  | 6586 |  | 6119 |  |
| Number of pseudogenes |  | 321 |  | 185 |  | NA |  |
| Mitochondrial genome |  |  |  |  |  |  |  |
| Assembly size (bases) |  | 5,991 |  | 5,989 |  | 5,994 |  |
| GC content (%) |  | 30.50 |  | 30.52 |  | 30.51 |  |
| Number of functional genes |  | 2 |  | 35 |  | 2 |  |
| Apicoplast genome |  |  |  |  |  |  |  |
| Assembly size (kb) |  | 26.60 |  | 29.58 |  | 34.53 |  |
| GC content (%) |  | 12.82 |  | 13.30 |  | 14.37 |  |
| Number of functional genes |  | 39 |  | 55 |  | 54 |  |
| Number of pseudogenes |  | 1 |  | 0 |  | NA |  |

The PvPAM genome contains 6497 functional genes (i.e., genes not annotated as a pseudogene), a higher number than found in PvW1, but still less than in PvP01 (Table 1). Out of the 6497 functional genes, 1109 are not present in PvP01. Of those, 487 genes cannot be linked to any PvP01 gene through orthology, and are therefore new in the PvPAM reference genome (Supplementary Table 1). The number of PvPAM pseudogenes is higher than what was observed for PvP01 (4.7% vs 2.7% of all genes). Pseudogenes are mainly located in the subtelomeres (89% of PvPAM and 95% of PvP01 pseudogenes), due to the dynamic nature of this region (Mok et al., 2008).

### PvPAM multigene families in subtelomeric or internally variable regions

Multigene families located at the subtelomeres or internally variable regions often play an important role in host-parasite interactions, immune response and pathogenicity, and can vary in (pseudo)gene number across isolates due to the dynamic nature of subtelomeres (Carlton et al., 2008). In particular, the *P. vivax* variant genes (*vir*) are known to be involved in immune invasion, and the *P. vivax* tryptophan-rich antigens (PvTRAG), *P. vivax* reticulocyte binding proteins (PvRBP) and *P. vivax* merozoite surface proteins (MSP) take part in invasion or red blood cell (RBC) binding.

In the PvPAM genome, 975+99 *vir* genes (functional genes + pseudogenes) were found, which is less than the 1093+116 in PvP01 and 1145 functional *vir* genes in PvW1. Remarkably, out of these 975 functional *vir* genes identified in PvPAM, only 370 have been transferred from the PvP01 reference genome through sequence similarity. The remaining 605 *vir* genes are newly predicted in PvPAM.

All 40 *PvTRAg* genes present in PvP01 were also found in PvPAM. No additional PvPAM *PvTRAg’s* were discovered based on othology groups (OrthoMCL) or the presence of the *Plasmodium* tryptophan-rich protein Pfam domain (PF12319). *PvTRAg36* (PVP01_0000140), which was reported to bind to the band 3 and basigin RBC receptors (Zeeshan et al., 2014;Alam et al., 2016), is annotated as a pseudogene in PvPAM. However, pseudogene annotation is based on prediction models. The utilization of an alternative translation start site and the splicing out of an intron (as seen in PvP01 PvTRAg36) would result in a fully functional PvTRAG36 protein. The more complete PvPAM assembly also enabled to locate 10 *PvTRAg’s* to chromosome 13, that were formerly annotated to the unassigned contigs ‘Transfer.PvP01_00_1.final’ (PvP01) and ‘CAJZCX010000017’ (PvW1).

Furthermore, the PvPAM genome contains all 5 full length *PvRBP* genes that are present in PvP01. Out of the 3 partial *PvRBP* genes that were identified in PvP01 and PvSalI, *PvRBP1-p1* (PVP01_0010770) and *PvRBP2-p2* (PVX_101590) are absent in PvPAM (Supplementary Table 2). *PvRBP2-p2* was reported to be present only in a subset of *P. vivax* samples, and is also absent in PvP01 (Hietanen et al., 2016). Of the 3 *PvRBP* pseudogenes in PvP01, *PvRBP3* and the isolate-specific *PvRBP2e* (Hester et al., 2013) are identified in PvPAM. The PvP01 *PvRBP2d* pseudogene is replaced by 3 partial genes in PvPAM, while in PvW1 it consists of 2 partial genes (alignment in Supplementary Figure 2, Supplementary Table 2). Further studies are required to confirm whether some isolates indeed show several shorter (partial) PvRBP2d proteins, or whether this is an annotation model artefact and in reality is one long pseudogene. No additional *PvRBP* genes were discovered in PvPAM based on orthology clusters or Pfam domains (PF16830, PF18515).

Next, the subtelomeric erythrocyte binding-like (EBL) superfamily contains two reticulocyte invasion-associated proteins in *P. vivax*. The *P. vivax* Duffy binding protein (PvDBP), the invasion ligand binding to the Duffy receptor (Barnwell et al., 1989), and PvDBP2, which shows reticulocyte binding activity (Ntumngia et al., 2016). *PvDBP* copy number variations have been associated with immune evasion (Hostetler et al., 2016;Popovici et al., 2020), but no additional *PvDBP* orthologs were identified in the PvPAM assembly. When PvPAM PacBio reads were mapped against the PvPAM genome, no increase in *PvDBP* or *PvDBP2* coverage (indicative of a duplication) was observed either (Supplementary Figure 3).

Additionally, multigene families that are located in tandem on hypervariable regions in the core genome, such as PvMSP3, PvMSP7 and the *P. vivax* serine-repeat antigens (PvSERA), have the potential to vary in number of genes (Pearson et al., 2016). The *PvMSP3* genes are highly polymorphic (Ford et al., 2020), and have been shown to have a varying number of genes at the center of the gene cluster (Minassian et al., 2021). Indeed, PvPAM contains two more centrally located *MSP3* genes (*MSP3.6, MSP3.7*) that are absent in PvP01 and PvW1, a feature it shares with the Central-American PvSalI. On the other hand, PvSalI’s *MSP3.4* is missing in PvPAM (Supplementary Figure 4). There was no variation in the number of genes observed in the *PvMSP7* and *PvSERA* families between PvPAM and PvP01.

### PvPAM drug resistance associated genes

Although drug resistance-associated mutations remain largely elusive in *P. vivax*, antifolate-resistance SNPs have been described for *dhps* and *dhfr* (Noviyanti et al., 2020). Antifolates are no longer recommended as a treatment in Peru, but the DHPS A383G and DHFR S58R and S117N resistance mutations are still highly prevalent in recently collected Peruvian samples (Flannery et al., 2015;Kattenberg et al., 2022;Villena et al., 2022). As expected, these mutations are also present in the PvPAM genome. The PvP01 reference contains the same antifolate-resistance mutations as PvPAM, while PvW1 possesses a different combination of antifolate resistance SNPs (Supplementary Table 3). Although the PvSalI genome is geographically the closest to PvPAM, PvSalI is still fully sensitive to antifolate drugs (Supplementary Table 3), as it was collected almost 20 years before the first resistant cases were described (Collins et al., 1972;Rieckmann et al., 1989). Overall, the PvPAM *dhps* and *dhfr* genes align well to their PvSalI, PvP01 and PvW1 homologs (≥98.3% and ≥99.8% sequence identity, respectively), except for an in-frame increase of tandem repeats upstream of the final *dhps* intron in PvPAM (Supplementary Figure 5A,B). Alignments of the chloroquine resistance-associated genes *mdr1* and *crt* (Noviyanti et al., 2020) of PvPAM, PvSalI, PvP01 and PvW1 show close similarities. PvPAM *mdr1* has 99.9% sequence identity to the *mdr1* genes in the 3 other reference genomes (Supplementary Figure 5C). PvPAM *crt* showed a lower sequence identity of 96.8%-99.2% to the *crt* sequence in the other reference genomes, due to PvPAM indels in introns 5, 9 and 12 that are caused by variations in tandem repeats (Supplementary Figure 5D).

Gene duplications have been associated with antimalarial resistance as well, and the PvPAM PacBio reads were mapped to the PvPAM reference genome to check for increased coverage indicative of a potential duplication. In addition, a search for new orthology group members of resistance-associated genes was done. In South America, duplications of *crt* (potential chloroquine resistance), *dhfr* and *dhfs* (potential antifolate resistance), and *mrp1* (potential atovaquone resistance) have been described (Flannery et al., 2015;Silva et al., 2018), but were not observed in PvPAM (Supplementary Figure 3). Mefloquine resistance-associated *mdr1* gene duplications have mainly been observed in Southeast Asia (Imwong et al., 2008;Suwanarusk et al., 2008;Lin et al., 2013;Auburn et al., 2016;Costa et al., 2017), and were not found in the Peruvian PvPAM genome (Supplementary Figure 3).

### South American samples mapped to PvPAM show more mapped reads and less read truncation

To investigate the difference in number of mapping reads between PvPAM, PvP01 and PvW1 as reference genome for South American WGS samples, WGS Illumina reads from 354 publicly available South American isolates and 63 non-South American isolates coming from Africa (N=18) and Asia (N=45), were mapped to the PvPAM, PvP01 and PvW1 reference genomes (overview of accession numbers and country of origin in Supplementary File 1).

When South American WGS reads are mapped to each of the three reference genomes, differences in number of mapped reads are small. 0.02%-0.09% more reads align to PvPAM than to PvW1 or PvP01 (Figure 2A; Wilcox *p* <0.01 and <0.0001 respectively). Additionally, 0.70% more non-South American reads are mapped to PvPAM than to PvP01 (Wilcox *p* <0.0001), but there is no difference in the number of mapped reads between PvPAM and PvW1 (Wilcox *p* non-significant; NS) (Figure 2A). The reference genome’s effect on read alignment becomes more apparent when chimaeric (truncated) reads are analysed separately. Mapping of South American reads to PvPAM results in 13.94%-41.77% less chimaeric reads compared to mapping to PvW1 or PvP01 (Figure 2B; Wilcox *p* <0.0001 for both). This indicates a higher structural similarity between South American isolates and PvPAM than with other reference genomes. Indeed, when reads from non-South American samples are mapped to the 3 reference genomes, the number of chimaeric reads is comparable between PvPAM and PvP01 (Wilcox *p* NS), and 0.24% lower in PvW1 (Wilcox *p* <0.05). This is in line with expectations, since 45 out of the 63 non-South American isolates are of Asian origin, and PvW1 originates from a Thai isolate that was sequenced with PacBio (Figure 2B).

**Figure 2.**
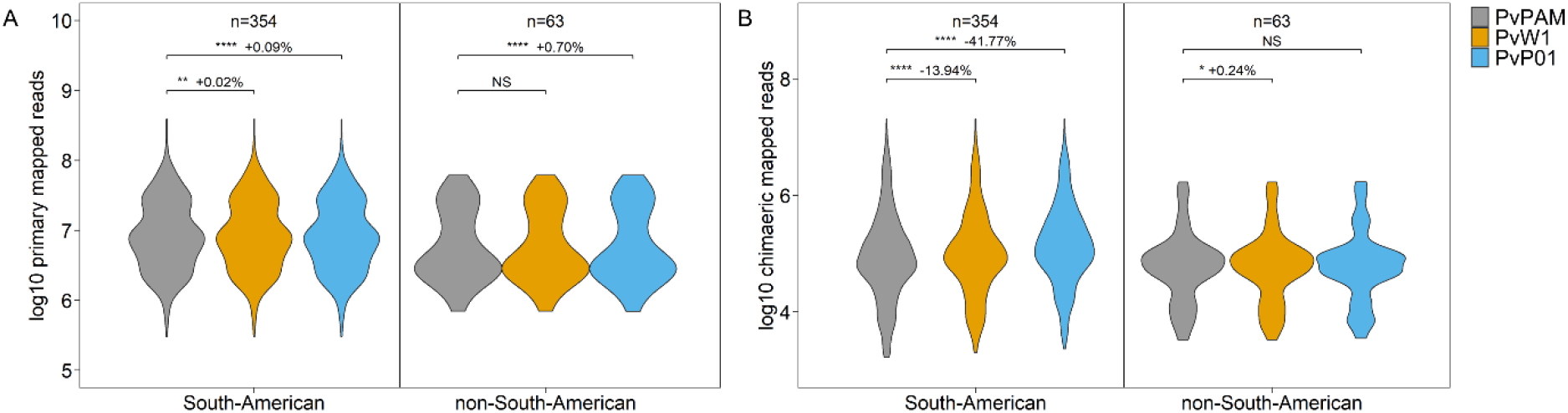
Violin plots showing the number of primary (**A**) and chimaeric (truncated; **B**) reads (log10 transformed) mapped to the PvPAM, PvW1 or PvP01 reference genomes. Reads originate from South American and non-South American Illumina WGS samples, which are shown separately. NS non-significant, * p<0.05, ** p<0.01, *** p<0.001, **** p<0.0001 (paired Wilcoxon signed-rank test). Next to the significance value, the % increase (+) or decrease (-) of mapped reads to PvPAM is shown. It is not possible to compare number of mapped reads between the South American and non-South American region, since those reads originate from different samples. n=number of Illumina WGS samples.

### PvPAM better reflects the South American variants

Since the PvPAM reference is derived from a Peruvian isolate, it likely resembles other South American isolates. Therefore, it is hypothesised that South American reads will contain less variants (SNPs, indels) when they are mapped to the PvPAM genome in comparison to PvP01 (Papua Indonesia) and PvW1 (Thailand). To investigate this, 354 publicly available Illumina WGS reads from South American samples (Peru, Brazil and Colombia; no other South American countries with >10 WGS isolates were found in our public data search) were mapped to the PvPAM, PvP01 and PvW1 reference genomes and variant called. The same was done for 63 Illumina WGS samples from Africa (Eritrea, Ethiopia, Sudan, Uganda), South Asia (Afghanistan, India, Pakistan) and Southeast Asia (Cambodia, Malaysia, Myanmar, Papua New Guinea, Thailand, Vietnam). Since PvPAM has a slightly longer subtelomeric genome than PvP01 and PvW1 (9.4 Mb vs 9.0 Mb) and since subtelomeric and core regions of the genome typically exhibit a different pattern of variant density (Supplementary Figure 6), variants of the core and subtelomeric genome were analysed separately.

When WGS reads from South American samples are mapped to the PvPAM genome, the number of variants in the core genome is significantly lower in comparison to PvP01 or PvW1 (Table 2, Supplementary Figure 7A,C). Interestingly, alignment of non-South American reads to the PvPAM genome still results in less core genome variants than when PvP01 is used as reference, but more core genome variants than with PvW1 as reference (Table 2, Supplementary Figure 7A,C). When the non-South American samples are divided into specific geographical regions, it becomes clear that reads from African and South Asian samples still contain the least core genome SNPs when mapped to PvPAM. Reads from Southeast Asian samples on the other hand, contain less core genome SNPs when mapped to PvP01 or PvW1 in comparison to PvPAM (Table 2, Supplementary Figure 7B). When indels are assessed, African reads still contain the least indels when mapped to PvPAM, but both South Asian as Southeast Asian reads now have the least indels when aligned to PvW1. In each case though, the use of PvP01 results in the highest number of indels, highlighting the advantage of structurally more correct PacBio references (PvW1, PvPAM) in the context of indels (Table 2, Supplementary Figure 7D). Overall, PvPAM is the most suited reference genome for use with South American and African WGS reads to investigate core genome variants, while PvW1 or PvP01 are a better choice for Southeast Asian reads. Both PvW1 and PvPAM are suited for South Asian reads, depending on whether core genome indels or SNPs are the main interest.

**Table 2.** Median number of variants depends on the used reference genome and region of origin. Numbers indicate the decrease (white background) or increase (grey background) in median number of variants per 10,000 bp when Illumina WGS reads are mapped to PvPAM in comparison to PvP01 or PvW1. For example, South American reads show 5.26 core genome SNPs less per 10,000 bp when mapped to the PvPAM genome in comparison to the PvP01 genome. Values are shown separately for the core genome and subtelomeric genome, and variants are divided into SNPs and indels (insertions and deletions). * p<0.05, ** p<0.01, *** p<0.001, **** p<0.0001 (paired Wilcoxon signed-rank test). African samples originate from Eritrea (5), Ethiopia (5), Sudan (5) and Uganda (3), South Asian samples from Afghanistan (5), India (5) and Pakistan (5), and Southeast Asian samples from Cambodia (5), Malaysia (5), Myanmar (5), Papua New Guinea (5), Thailand (5) and Vietnam (5). N=number of Illumina WGS samples.

|  | PvPAM vs PvP01 |  |  |  | PvPAM vs PvW1 |  |  |  | N |
| --- | --- | --- | --- | --- | --- | --- | --- | --- | --- |
|  | SNPs |  | indels |  | SNPs |  | indels |  |  |
| CORE GENOME |  |  |  |  |  |  |  |  |  |
| South America | 5.26 | **** | 2.41 | **** | 5.34 | **** | 2.10 | **** | 365 |
| Peru | 5.61 | **** | 2.47 | **** | 5.63 | **** | 2.17 | **** | 137 |
| Brazil | 4.82 | **** | 2.40 | **** | 4.90 | **** | 2.10 | **** | 152 |
| Colombia | 4.40 | **** | 2.31 | **** | 4.56 | **** | 2.00 | **** | 65 |
| Non-South America | 0.98 | * | 0.94 | **** | 0.63 | * | 0.03 | **** | 63 |
| Africa | 2.15 | **** | 0.96 | **** | 0.80 | **** | 0.11 | ** | 18 |
| South Asia | 2.24 | **** | 1.03 | **** | 0.57 | **** | 0.04 | * | 15 |
| Southeast Asia | 0.39 | **** | 0.25 | **** | 2.08 | **** | 0.66 | **** | 30 |
| SUBTELOMERIC GENOME |  |  |  |  |  |  |  |  |  |
| South America | 4.55 | **** | 0.45 | **** | 5.21 | **** | 0.48 | **** | 365 |
| Peru | 1.63 | **** | 0.15 | **** | 1.67 | **** | 0.12 | **** | 137 |
| Brazil | 4.18 | **** | 0.30 | **** | 5.09 | **** | 0.43 | **** | 152 |
| Colombia | 5.18 | **** | 0.67 | **** | 7.41 | **** | 0.85 | **** | 65 |
| Non-South America | 2.63 | **** | 0.18 | **** | 1.41 | **** | 0.05 | **** | 63 |
| Africa | 1.61 | **** | 0.23 | **** | 0.82 | **** | 0.14 | *** | 18 |
| South Asia | 1.63 | **** | 0.18 | **** | 0.42 | *** | 0.06 | * | 15 |
| Southeast Asia | 3.96 | **** | 0.34 | **** | 3.01 | **** | 0.16 | **** | 30 |

In the subtelomeric genome, reads from South American samples again show lower numbers of variants when mapped to PvPAM in comparison to the other 2 genomes (Table 2, Supplementary Figure 8A-D). Reads from non-South American samples, however, now also show less subtelomeric variants when mapped to PvPAM than when mapped to PvP01 or PvW1 (Table 2, Supplementary Figure 8A,C), and this is true for all the different geographical regions assessed (Table 2, Supplementary Figure 8B,D). This shows that the PvPAM subtelomeres are more similar to the subtelomeric regions of isolates from all continents than the PvP01 or PvW1 subtelomeres, although the PvPAM subtelomeres still resemble the South American subtelomeres the most (largest decrease in median number of variants – Table 2).

It can be concluded that the PvPAM reference reflects South American variants the best, and therefore, the generated vcf file containing variants of South American reads mapped to the PvPAM reference genome, can be of use for future research (available on the European Variation Archive (EVA)).

## Discussion

This study presents the first high-quality South American *P. vivax* reference genome, named PvPAM. Although South American *P. vivax* isolates are genetically distinct from the other continents (Benavente et al., 2021) and are the predominant cause of malaria in South America (WHO, 2021), no qualitative South American *P. vivax* reference genome existed until now. When compared to PvW1 and PvP01, PvPAM is the best suited reference genome for alignment of South American WGS reads. PvPAM was derived from a monoclonal *P. vivax* malaria patient from the Peruvian Amazon, and the isolate did not undergo culture adaptation prior to sequencing. Only 9 ng of DNA extracted from 100 μl of leukocyte depleted RBCs (1% parasitaemia) was used for PacBio sequencing, in contrast to the 107 ng obtained from controlled human infections for the construction of the PvW1 PacBio reference genome (Minassian et al., 2021).

To assess the benefit of using a South American reference for read alignment of South American samples, PvPAM was compared to PvW1 and PvP01, the two other high-quality reference genomes existing for *P. vivax*. A small but significantly higher number of South American reads mapped to PvPAM in comparison to these other two references. More importantly, 14% and 42% less truncated reads were found compared to PvW1 and PvP01, respectively. For non-South-American samples, alignment to PvW1 and PvPAM resulted in a similarly high number of mapped reads, but use of the PacBio-based Thai PvW1 reference resulted in less truncated reads. Since read truncation is typically due to large structural variants (e.g. indels >50 bp, translocations, inversions, copy number variations), this points towards continent-specific structural differences between the used reference genomes. In terms of number of mapped and truncated reads, PvPAM is well suited as a reference genome for South American WGS data, and is competitive with PvP01 and PvW1 for general use with WGS data from other continents.

The advantage of using a reference genome that is closely related to the genome of the sequenced isolate was further demonstrated by investigating the number of core genome variants. Alignment of South American WGS reads to the PvPAM reference showed the lowest number of core genome variants, indicating that South American samples resemble PvPAM most closely. Similarly, PvW1, originating from a Thai isolate, is the best reference to minimize the number of core genome variants in Southeast Asian samples. Alignment of African WGS reads to PvPAM still resulted in the lowest number of core genome variants, while alignment of South Asian reads to PvPAM and PvW1 produced the lowest number of core genome SNPs and indels, respectively. These findings mirror phylogenetic trees in which South American isolates are genetically most distinct from Southeast Asian isolates, but more similar to African and, to a lesser extent, South Asian isolates (Benavente et al., 2021).

Alignment of WGS reads to PvPAM always resulted in the lowest number of subtelomeric variants, regardless of the sample’s country of origin. Interestingly, even Southeast Asian samples showed less subtelomeric variants when mapped to PvPAM compared to the Thai and Pacbio-based PvW1. However, *Plasmodium* subtelomeric regions remain largely elusive, and the extent to which the subtelomeric sequences vary between isolates and show geographical clustering is still uncertain. The subtelomeric regions of *P. falciparum* have been shown to vary in length across isolates (Otto et al., 2018), and similarly, PvPAM has longer subtelomeric regions than PvP01 or PvW1. The longer subtelomeres of PvPAM contain unique regions that are not present in either PvP01 or PvW1, which might result in a subtelomeric backbone that is better suited to align samples from all continents against. This could improve accuracy of read mapping and thus lead to lower variant densities.

The high-quality and longer subtelomeric regions of the PvPAM genome provide new insights into the subtelomeric gene families and their diversity. A comparison of *vir* genes between PvP01 and PvPAM reveals that both have a high number of unique functional *vir* genes that are absent in the other genome (723 and 605, respectively). This could be due to geographical differences between PvP01 and PvPAM, but it is also possible that there is an inherent variability in *vir* genes, even across isolates from the same region. Other subtelomeric gene families did not show a gain or loss of gene members in PvPAM and appear to be more stable than the *vir* family. However, more *P. vivax* assemblies with high-quality subtelomeres are needed to study variation of *vir* genes and other subtelomeric multigene families in depth. Some subtelomeric genes, such as the *PvTRAg25-PvTRAg33* cluster, could for the first time be located to a chromosome. On the other hand, 12 subtelomeric PvPAM contigs could not be assigned to any chromosome based on PvP01 or PvW1 sequence similarity.

In this study, the core genome of PvPAM as well as PvP01 and PvW1 was defined based on GC-content and pairwise alignment between these three genomes. Although no standard definition is available for subtelomeric-core genome boundaries, GC-content has been used earlier as a marker for *P. vivax* subtelomeres (Carlton et al., 2008). The provided core genome regions could be further refined by comparing positionally conserved genes across different *P. vivax* isolates, as was done for *P. falciparum* isolates (Otto et al., 2018). This, again, would require multiple high-quality *P. vivax* subtelomere assemblies that are currently not available. However, the here used combination of long-read sequencing and a low-input material approach, might facilitate assembly of *P. vivax* field isolates from blood volumes as small as a finger prick. In addition, Oxford Nanopore’s portable sequencer could become a quick and easy to use alternative for PacBio sequencing (De Meulenaere et al., 2022a), given further improvements in accuracy and DNA input requirements.

In summary, the PvPAM reference genome is the best choice to align South American WGS data to, highlighting the importance of using a continent-specific reference. Furthermore, it performed well for African and, to a lesser extent, South Asian WGS data. When studying subtelomeric regions, PvPAM proved to be the most suited reference genome, regardless of the sample’s continent of origin. Overall, the PvPAM reference genome will improve the quality of genomic analyses conducted on South American samples or subtelomeric genomes.

## Methods

### Collection and processing of *P. vivax* patient blood samples

*P. vivax–*infected blood was collected from adult patients (≥ 18 years) with acute *P. vivax* infection from the Peruvian city of Iquitos and neighbouring communities. All *P. vivax* cases were diagnosed by light microscopy. 10-20 mL blood samples were collected in lithium-heparin tubes from patients with single *P. vivax* infections with parasite densities >0.1% and gametocyte proportion <50%.

Blood samples from patients with *P. vivax* infection were processed within 6 hours of collection. Leukocytes and platelets were depleted using cellulose columns (Sriprawat et al., 2009). Then, 100 μl of *P. vivax–*infected RBCs were stored at 50% haematocrit for future short- and long-read whole genome sequencing (WGS). The remaining blood was used for other purposes.

### DNA extraction and size selection of Pv01-19

Out of the 22 collected samples, sample Pv01-19 contained the highest parasitaemia of 1.0% (approximately 90% ring stages), and was therefore selected for long and short-read WGS. DNA was extracted from the stored 100 μl leukocyte-depleted RBCs using the QIAamp DNA mini kit (Qiagen), following the manufacturer’s instructions. After elution in 50 μL AE buffer, 0.46 ng/μl DNA was obtained. Fragments <3kb were removed with Ampure PB beads (2.2V of 40% beads), and DNA was eluted in 50 μl PacBio elution buffer, resulting in 0.21 ng/μl DNA. Input fragment target size for the PacBio ultralow input protocol is 8-10 kilobases (kb). Since the concentration of Pv01-19 was too low (<1 ng/μl), no Fragment Analyzer (Agilent) profile could be made. However, Fragment Analyzer profiles of higher concentration samples that were similarly stored fell within the 8-10 kb range, which is why no further shearing was applied on the Pv01-19 DNA.

### PacBio sequencing of Pv01-19

9 ng of DNA was used for library preparation with the SMRTbell Express Template Prep Kit 2.0, following the ultralow input protocol (version 01, August 2020). This protocol includes a 13-cycle PCR step to amplify the low amount of input DNA, using reagents from the SMRTbell gDNA Sample Amplification Kit (master mixes A and B cover different GC compositions). For the mix A PCR, 3 additional cycles were done to increase the yield. The final library was size-selected for fragments >6,800 bases with a BluePippin device (Sage Science), since the Fragment Analyzer profile showed a peak at 8,304 bases. After clean-up, a 14.3 ng/μl library was obtained, with a fragment length peak at 8,341 bases (Femto Pulse, Agilent). The Femto Pulse trace showed no signs of dimer or overamplification.

SMRTbell templates were annealed with Sequencing Primer v4 and bound to a polymerase using the Sequel II binding Kit 2.0. SMRT Link settings for sequencing were entered according to the ultralow input protocol (CCS mode, Iso-Seq Experiment enabled, Iso-Seq Version Express to enable ProNex Cleanup workflow, with ProNex cleanup of 50% anticipated). Next, the library was loaded onto an 8M SMRT cell and sequenced on a Sequel II with 2 hours pe-extension time and 30 hours movie time (Earlham Institute, Norwich). 6.1 million continuous long reads (CLR) were generated, resulting in a total number of 497 billion sequenced bases.

### Illumina sequencing of Pv01-19

To be able to polish PacBio sequencing errors, 1 ng DNA of the Pv01-19 sample was sent for Illumina sequencing (GenoScreen, Lille). The library was prepared with the Nextera XT DNA Sample Prep Kit (Illumina) following the manufacturer’s protocol, and sequenced on an Illumina NovaSeq 6000 platform with 2×150 bp paired-end reads. 68.3 million raw reads were generated, resulting in 10.3 billion sequenced bases.

### Pre-processing of PacBio reads

Reads were pre-processed with the PacBio secondary analysis tools available on PacBio Bioconda (https://github.com/PacificBiosciences/pbbioconda). Raw CLR subreads were converted to HiFi circular consensus sequencing (CCS) reads with pbccs v6.0.0 (--hifi-kinetics). During this process, 43% of the reads were lost, mainly due to misalignment of subreads from the same CLR read. To retain the ‘pw’ tag necessary for future Arrow assembly polishing, the ccs-kinetics-bystrandify tool (v2.0.0) from pbbam was used. PCR-adapters were trimmed with lima (--same --ccs --min-score 80), during which 5.6% of the CCS reads were lost. Next, duplicate removal with pbmarkdup v1.0.2 (--rmdup) caused another 5.6% of trimmed CCS read loss. This resulted in a final number of 24.5 billion CCS bases, with an average read length of 10564 bases. To remove human reads, reads were mapped to the human genome (GRCh38) with the minimap2 wrapper pbmm2 v1.8.0 (--preset CCS --sort -- unmapped), after which the unmapped reads were selected with samtools view (-f 4). Human reads represented only 0.18% of all CCS trimmed and non-duplicate bases.

Presence of contaminating species was checked with a Kraken2 run (--confidence 0.1; v2.0.9-beta) (Wood et al., 2019), during which the pre-processed and human-filtered reads were taxonomically classified with the PlusPF (09/08/22) database. No contaminants were identified that represented ≥0.01% of the reads.

Next, the pre-processed reads were mapped to the PvP01 reference genome (PlasmoDB, v51) with pbmm2 v1.8.0 to estimate the depth and check the similarity with the PvP01 reference genome.

### Assembly of the PvPAM reference genome

The pre-processed PacBio reads were *de novo* assembled into a draft genome with Canu v2.0 (-pacbio hifi) (Nurk et al., 2020). Next, contigs were scaffolded onto the PvP01 (PlasmoDB, v51) and PvW1 (PlasmoDB, v60) chromosomes using RagTag v2.1.0 (scaffold command, no prior correction done to prevent loss of strain-specific information) (Alonge et al., 2021), and additional verification with minimap2 (-ax map-pb). When a contig was scaffolded differently on PvP01 as compared to PvW1, it was manually checked to which chromosomal location the contig fit best or if it had to be removed from the scaffold due to low confidence. 12 contigs could not be scaffolded confidently to any chromosome and remained unassigned. No contigs were scaffolded to the mitochondrial chromosome, due to its small size of approx. 6 kb (smaller than mean read length). Therefore, CCS reads that aligned (pbmm2) to the PvP01 or PvW1 mitochondrial chromosome were selected, duplicate removed (pbmarkdup), and assembled separately with Canu. Duplicated regions (circular chromosome) were manually removed.

Gaps between contigs of the same scaffold were then manually patched based on the PvW1 and PvP01 reference genome, using Geneious Prime (2022 release). PvW1 was preferentially used for patching, except when the contig ends only aligned to PvP01 (only the case for one patch on chromosome 2). Dotplots were made to check for large duplications due to assembly errors or due the circular mitochondrial and apicoplast chromosomes, but none were observed.

The assembly was polished based on the pre-processed CCS reads with 3 Arrow iterations (GCpp v2.0.2, https://github.com/PacificBiosciences/gcpp), introducing 5,967 changes to the assembly. This was followed by three iterations of Pilon (v1.23) short read polishing, resulting in 970 more changes. For each round of Pilon polishing, Illumina reads were mapped to the assembly with bwa mem v0.7.17, sorted on coordinates (samtools sort), duplicate reads were removed with Picard’s MarkDuplicates v2.22.0 (REMOVE_DUPLICATES=true), and confidently mapped reads (mapping quality = 60, no secondary reads) were selected with samtools view (-q60 -F256). The latter is done to prevent the shorter Illumina reads from misaligning to repetitive regions and incorrect polishing.

Completeness of the assembly was checked with Busco v5.4.4 (Manni et al., 2021). Number of single-copy, fragmented and missing BUSCO’s (Benchmarking Universal Single-Copy Orthologs) are within the same range for PvPAM, PvP01 and PvW1 (Supplementary Table 4). Finally, the PvPAM assembly was annotated with Companion v1.0.2 (Steinbiss et al., 2016). Contiguation was turned off, reference proteins were aligned to the target sequence, and RATT was used for gene transfer (on species level) from the PvP01 reference genome. For other parameters, default settings were used.

### Defining the subtelomeric-core genome boundaries

Boundaries between the core genome and subtelomeric regions were here defined based on GC content. GC content was calculated per 1 kb window (sliding at steps of 100 bases) with bedtools v2.29.2 (makewindows, nuc). In R v4.2.1, draft subtelomeric boundaries were defined from both sides of the chromosome as the first window of a 10 kb region (100 consecutive windows) for which GC rose consistently above 35%. These draft core genome regions were then extracted from the PvPAM assembly with samtools faidx. Per chromosome, the PvPAM draft core region was aligned to the corresponding PvP01 (PlasmoDB, v46) and PvW1 chromosomes with Progressive Mauve (default settings) in Geneious Prime (Darling et al., 2010). The part of the draft core region that aligned to the 2 other reference genomes was considered to be the final core genome region, and these regions are shown in Supplementary Table 5.

Unassigned contigs were categorised as subtelomeric or core based on their GC content. If the GC content was below the mean GC content of the core and subtelomeric regions of the 14 chromosomes (as determined above), the contig was considered subtelomeric. This was the case for all contigs of PvPAM and PvW1, and for 207/226 contigs of PvP01 (Supplementary Table 5).

### Mapping of publicly available WGS data to PvPAM, PvP01 and PvW1 and variant calling

To be able to compare mapping quality and variants between the PvPAM, PvP01 and PvW1 genomes, and to construct a South American vcf (variant call format) file, 354 South American (152 Brazil, 65 Colombia, 137 Peru) and 63 non-South American (5 Afghanistan, 5 Cambodia, 5 Eritrea, 5 Ethiopia, 5 India, 5 Malaysia, 5 Myanmar, 5 Pakistan, 5 Papua New Guinea, 5 Sudan, 5 Thailand, 3 Uganda, 5 Vietnam) Illumina WGS samples were obtained from public databases (Dharia et al., 2010;Chan et al., 2012;Neafsey et al., 2012;Flannery et al., 2015;Delgado-Ratto et al., 2016;Hupalo et al., 2016;Pearson et al., 2016;de Oliveira et al., 2017;Cowell et al., 2018;de Oliveira et al., 2020;Benavente et al., 2021;Adam et al., 2022;De Meulenaere et al., 2022b;Kattenberg et al., 2022) and newly sequenced samples from Peru and Brazil (SRA BioProjects PRJNA853729 and PRJNA934307, respectively). Supplementary File 1 gives an overview of the used accession numbers, their country and the study they originate from.

Raw reads were mapped to the human reference genome (GRCh38) with bwa mem v0.7.17 and coordinate sorted with samtools, after which the non-human reads that did not map as a pair were selected (samtools view -F 2) and converted to the fastq format again (samtools fastq). Those were then mapped to the PvPAM, PvP01 (version 46) or PvW1 reference genome with bwa mem, coordinate sorted, and duplicate marked with Picard MarkDuplicates v2.22.0. Per sample, the median alignment score was extracted using an awk loop, and number of primary and chimaeric reads were extracted with samtools view -c (-F 256 -F 2048 -F 1024 and -f 2048 -F 1024 respectively). The same was done on only the core genome of those same mapped files, as defined in previous section (Supplementary Table 5).

Next variants were called using the GATK (v4.1.4.1) HaplotypeCaller (-ERC GVCF), and resulting gvcf files samples were combined (GATK CombineGVCFs) and converted (GATK GenotypeGVCFs) to a South American and a non-South American vcf file. Each vcf was split into a SNP and indel vcf (GATK SelectVariants), after which filter fields were defined according to the GATK golden standard with VariantFiltration (for the SNP vcf: QD<2.0, QUAL<30.0, SOR>3.0, FS>60.0, MQ<40.0, MQRankSum<-12.5, ReadPosRankSum<-8.0; for the indel vcf: QD<2.0, QUAL<30.0, FS>200.0, ReadPosRankSum<-20.0). Only variants that passed the filters described and did not have >50% of missing genotypes (bcftools view -i ‘F_MISSING<0.5’) were retained in the vcf files. Variants were annotated with SnpEff v4.3t and the -stats summary was used to compare the variant rate per base, mean allele frequency and mean indel length between the three used reference genomes. This was repeated on the same vcf files for which first the core genome was extracted (core genome regions in Supplementary Table 5). Vcf files are made available under Study PRJEB60575 on the European Nucleotide Archive, and Supplementary File 1 links the WGS samples names used in the vcf to their accession numbers.

### Multiplicity of infection (MOI) estimation of Pv01-19

To estimate the MOI of Pv01-19, its Illumina reads were mapped to PvP01 (PlasmoDB v46) with bwa mem v0.7.17 and coordinate sorted (samtools sort), and duplicate reads were flagged with Picard’s MarkDuplicates v2.22.0. The same was done for Illumina reads from 15 other Peruvian samples collected at the same location and in the same way (BioProject PRJNA853729) (De Meulenaere et al., 2022b), to provide samples for comparison to the MOI estimation tools. Pv01-19 was variant called together with the 15 other Peruvian samples following the same pipeline as described in previous section for the public data samples. Core and non-hypervariable regions as defined by Pearson et al. (2016) were extracted from the SNP vcf with bcftools tabix, after which estMOI v1.03 (Assefa et al., 2014) and the moimix R package (Lee and Bahlo, 2016) were ran. Both tools estimated a MOI=1 for Pv01-19.

### Statistical analysis

Non-parametrical paired Wilcoxon signed-rank tests were performed in R v4.2.1 (R Core Team, 2019).

## Supporting information

Supplementary File 2 (assembly)

Supplementary File 3 (annotation)

## Declarations

### Ethics approval and consent to participate

Ethical approval for the collection and sequencing of *P. vivax* isolates from adult patients in Iquitos (Peru) was obtained from the Institute of Tropical Medicine Antwerp (ITM) Institutional Review Board (IRB; protocol 1345/19) and the ethics committee at the University Hospital of Antwerp (protocol B3002020000016 and B300201523588) and Universidad Peruana Cayetano Heredia (UPCH; Lima, Peru) (protocol 101898).

All participants provided written informed consent before enrolment. The study was conducted according to the principles stated in the Declaration of Helsinki 2013 (World Medical Association, 2013).

### Availability of data and materials

The annotated reference genome and raw PacBio and Illumina reads used for its construction are available under Study PRJEB59758 in the European Nucleotide Archive (https://www.ebi.ac.uk/ena/). Additionally, the reference genome (.fasta.gz) and annotation file (gff3 format) are made available in Supplementary Files 2 and 3, respectively. Newly generated WGS Illumina reads from Peruvian (N=7) and Brazilian (N=21) samples are made available under BioProjects PRJNA853729 and PRJNA934307 in the Sequence Read Archive, respectively (https://www.ncbi.nlm.nih.gov/sra). The variant files (.vcf) containing SNPs and indels of publicly available South American WGS Illumina data mapped to the PvPAM genome, are available under Study PRJEB60575 in the European Nucleotide Archive (https://www.ebi.ac.uk/ena/). Supplementary File 1 links the sample names used in the vcf to their accession numbers.

### Authors’ contributions

ARU, KDM, BC and KL conceived the project. ARU supervised the overall study, KL and BC supervised the bioinformatic work, and DG supervised the field collection. KDM designed and carried out the wet lab work, led the bioinformatic analysis and interpreted the results with contributions from BC and ARU. KDM, BC and ARU wrote the first draft of the manuscript. All authors read and approved the final manuscript.

## Acknowledgements

We would like to thank the patients, clinical staff, microscopists, and field staff from health centers and hospitals in and around Iquitos (Peru) for their cooperation and contribution to the collection of *P. vivax* isolates. In particular, Elizabeth Villasis (Universidad Peruana Cayetano Heredia), for the organisation and support of the sample collection at Amazonian-ICEMR laboratory facilities in Iquitos. We are grateful to Prof. Marcelo Urbano Ferreira (University of São Paulo, Brazil) for his contribution of WGS Illumina data from 21 Brazilian samples, and Dr. Pieter Monsieurs (Institute of Tropical Medicine Antwerp) for his help with the collection of publicly available WGS Illumina data from South American samples.

## Supplementary material

**Supplementary Data.pdf**

Contains Supplementary Figures 1-8 and Supplementary Tables 1-4.

**Supplementary Table 5.xlsx**

Overview of the core genome regions of PvPAM, PvP01 and PvW1, as defined in the Methods section. Per chromosome or unassigned contig, the start and end position of the core genome is given.

**Supplementary File 1.xlsx**

Accession numbers of the 354 publicly available WGS Illumina reads from South American isolates, and 63 non-South American isolates coming from Africa and South Asia and Southeast Asia, which were used to map against the PvPAM, PvP01 and PvW1 reference genomes.

**Supplementary File 2.fasta.gz**

The PvPAM sequence in fasta format (gzipped).

**Supplementary File 3.txt**

Annotation of the PvPAM reference genome in gff3 format.

## References

Adam, I., Alam, M.S., Alemu, S., Amaratunga, C., Amato, R., Andrianaranjaka, V., Anstey, N.M., Aseffa, A., Ashley, E., and Assefa, A. (2022). An open dataset of Plasmodium vivax genome variation in 1,895 worldwide samples. Wellcome Open Research 7.

Alam, M.S., Zeeshan, M., Rathore, S., and Sharma, Y.D. (2016). Multiple Plasmodium vivax proteins of Pv-fam-a family interact with human erythrocyte receptor Band 3 and have a role in red cell invasion. Biochemical and biophysical research communications 478, 1211–1216.

Alonge, M., Lebeigle, L., Kirsche, M., Aganezov, S., Wang, X., Lippman, Z.B., Schatz, M.C., and Soyk, S. (2021). Automated assembly scaffolding elevates a new tomato system for high-throughput genome editing. BioRxiv.

Assefa, S.A., Preston, M.D., Campino, S., Ocholla, H., Sutherland, C.J., and Clark, T.G. (2014). estMOI: estimating multiplicity of infection using parasite deep sequencing data. Bioinformatics 30, 1292–1294.

Auburn, S., Böhme, U., Steinbiss, S., Trimarsanto, H., Hostetler, J., Sanders, M., Gao, Q., Nosten, F., Newbold, C.I., and Berriman, M. (2016). A new Plasmodium vivax reference sequence with improved assembly of the subtelomeres reveals an abundance of pir genes. Wellcome open research 1.

Barnwell, J.W., Nichols, M.E., and Rubinstein, P. (1989). In vitro evaluation of the role of the Duffy blood group in erythrocyte invasion by Plasmodium vivax. The Journal of experimental medicine 169, 1795–1802.

Benavente, E.D., Manko, E., Phelan, J., Campos, M., Nolder, D., Fernandez, D., Velez-Tobon, G., Castaño, A.T., Dombrowski, J.G., and Marinho, C.R. (2021). Distinctive genetic structure and selection patterns in Plasmodium vivax from South Asia and East Africa. Nature communications 12, 1–11.

Bermúdez, M., Moreno-Pérez, D.A., Arévalo-Pinzón, G., Curtidor, H., and Patarroyo, M.A. (2018). Plasmodium vivax in vitro continuous culture: the spoke in the wheel. Malaria journal 17, 301.

Carlton, J.M., Adams, J.H., Silva, J.C., Bidwell, S.L., Lorenzi, H., Caler, E., Crabtree, J., Angiuoli, S.V., Merino, E.F., and Amedeo, P. (2008). Comparative genomics of the neglected human malaria parasite Plasmodium vivax. Nature 455, 757.

Chan, E.R., Menard, D., David, P.H., Ratsimbasoa, A., Kim, S., Chim, P., Do, C., Witkowski, B., Mercereau-Puijalon, O., and Zimmerman, P.A. (2012). Whole genome sequencing of field isolates provides robust characterization of genetic diversity in Plasmodium vivax. PLoS neglected tropical diseases 6, e1811.

Collins, W.E., Contacos, P.G., Krotoski, W.A., and Howard, W.A. (1972). Transmission of four Central American strains of Plasmodium vivax from monkey to man. The Journal of Parasitology, 332–335.

Costa, G.L., Amaral, L.C., Fontes, C.J.F., Carvalho, L.H., De Brito, C.F.A., and De Sousa, T.N. (2017). Assessment of copy number variation in genes related to drug resistance in Plasmodium vivax and Plasmodium falciparum isolates from the Brazilian Amazon and a systematic review of the literature. Malaria journal 16, 1–11.

Cowell, A.N., Valdivia, H.O., Bishop, D.K., and Winzeler, E.A. (2018). Exploration of Plasmodium vivax transmission dynamics and recurrent infections in the Peruvian Amazon using whole genome sequencing. Genome medicine 10, 52.

Darling, A.E., Mau, B., and Perna, N.T. (2010). progressiveMauve: multiple genome alignment with gene gain, loss and rearrangement. PloS one 5, e11147.

De Meulenaere, K., Cuypers, W.L., Rosanas-Urgell, A., Laukens, K., and Cuypers, B. (2022a). Selective whole-genome sequencing of Plasmodium parasites directly from blood samples by Nanopore adaptive sampling. bioRxiv, 2022.2011. 2029.518068.

De Meulenaere, K., Prajapati, S.K., Villasis, E., Cuypers, B., Kattenberg, J.H., Kasian, B., Laman, M., Robinson, L.J., Gamboa, D., and Laukens, K. (2022b). Band 3–mediated Plasmodium vivax invasion is associated with transcriptional variation in PvTRAg genes. Frontiers in Cellular and Infection Microbiology, 1460.

De Oliveira, T.C., Corder, R.M., Early, A., Rodrigues, P.T., Ladeia-Andrade, S., Alves, J.M.P., Neafsey, D.E., and Ferreira, M.U. (2020). Population genomics reveals the expansion of highly inbred Plasmodium vivax lineages in the main malaria hotspot of Brazil. PLoS neglected tropical diseases 14, e0008808.

De Oliveira, T.C., Rodrigues, P.T., Menezes, M.J., Goncalves-Lopes, R.M., Bastos, M.S., Lima, N.F., Barbosa, S., Gerber, A.L., Loss De Morais, G., and Berna, L. (2017). Genome-wide diversity and differentiation in New World populations of the human malaria parasite Plasmodium vivax. PLoS neglected tropical diseases 11, e0005824.

Delgado-Ratto, C., Gamboa, D., Soto-Calle, V.E., Van Den Eede, P., Torres, E., Sánchez-Martínez, L., Contreras-Mancilla, J., Rosanas-Urgell, A., Rodriguez Ferrucci, H., and Llanos-Cuentas, A. (2016). Population genetics of Plasmodium vivax in the Peruvian Amazon. PLoS neglected tropical diseases 10, e0004376.

Dharia, N.V., Bright, A.T., Westenberger, S.J., Barnes, S.W., Batalov, S., Kuhen, K., Borboa, R., Federe, G.C., Mcclean, C.M., and Vinetz, J.M. (2010). Whole-genome sequencing and microarray analysis of ex vivo Plasmodium vivax reveal selective pressure on putative drug resistance genes. Proceedings of the National Academy of Sciences, 201003776.

Flannery, E.L., Wang, T., Akbari, A., Corey, V.C., Gunawan, F., Bright, A.T., Abraham, M., Sanchez, J.F., Santolalla, M.L., and Baldeviano, G.C. (2015). Next-generation sequencing of Plasmodium vivax patient samples shows evidence of direct evolution in drug-resistance genes. ACS infectious diseases 1, 367–379.

Fola, A.A., Harrison, G.A., Hazairin, M.H., Barnadas, C., Hetzel, M.W., Iga, J., Siba, P.M., Mueller, I., and Barry, A.E. (2017). Higher complexity of infection and genetic diversity of Plasmodium vivax than Plasmodium falciparum across all malaria transmission zones of Papua New Guinea. The American journal of tropical medicine and hygiene 96, 630.

Ford, A., Kepple, D., Abagero, B.R., Connors, J., Pearson, R., Auburn, S., Getachew, S., Ford, C., Gunalan, K., and Miller, L.H. (2020). Whole genome sequencing of Plasmodium vivax isolates reveals frequent sequence and structural polymorphisms in erythrocyte binding genes. PLoS neglected tropical diseases 14, e0008234.

Gardner, M.J., Hall, N., Fung, E., White, O., Berriman, M., Hyman, R.W., Carlton, J.M., Pain, A., Nelson, K.E., and Bowman, S. (2002). Genome sequence of the human malaria parasite Plasmodium falciparum. Nature 419, 498–511.

Hester, J., Chan, E.R., Menard, D., Mercereau-Puijalon, O., Barnwell, J., Zimmerman, P.A., and Serre, D. (2013). De novo assembly of a field isolate genome reveals novel Plasmodium vivax erythrocyte invasion genes. PLoS neglected tropical diseases 7, e2569.

Hietanen, J., Chim-Ong, A., Chiramanewong, T., Gruszczyk, J., Roobsoong, W., Tham, W.-H., Sattabongkot, J., and Nguitragool, W. (2016). Gene models, expression repertoire, and immune response of Plasmodium vivax reticulocyte binding proteins. Infection and immunity 84, 677–685.

Hostetler, J.B., Lo, E., Kanjee, U., Amaratunga, C., Suon, S., Sreng, S., Mao, S., Yewhalaw, D., Mascarenhas, A., and Kwiatkowski, D.P. (2016). Independent origin and global distribution of distinct Plasmodium vivax Duffy binding protein gene duplications. PLoS neglected tropical diseases 10, e0005091.

Howes, R.E., Battle, K.E., Mendis, K.N., Smith, D.L., Cibulskis, R.E., Baird, J.K., and Hay, S.I. (2016). Global epidemiology of Plasmodium vivax. The American journal of tropical medicine and hygiene 95, 15.

Hupalo, D.N., Luo, Z., Melnikov, A., Sutton, P.L., Rogov, P., Escalante, A., Vallejo, A.F., Herrera, S., Arévalo-Herrera, M., and Fan, Q. (2016). Population genomics studies identify signatures of global dispersal and drug resistance in Plasmodium vivax. Nature genetics 48, 953–958.

Imwong, M., Pukrittayakamee, S., Pongtavornpinyo, W., Nakeesathit, S., Nair, S., Newton, P., Nosten, F., Anderson, T.J., Dondorp, A., and Day, N.P. (2008). Gene amplification of the multidrug resistance 1 gene of Plasmodium vivax isolates from Thailand, Laos, and Myanmar. Antimicrobial agents and chemotherapy 52, 2657–2659.

Karst, S.M., Ziels, R.M., Kirkegaard, R.H., Sørensen, E.A., Mcdonald, D., Zhu, Q., Knight, R., and Albertsen, M. (2021). High-accuracy long-read amplicon sequences using unique molecular identifiers with Nanopore or PacBio sequencing. Nature Methods 18, 165–169.

Kattenberg, J.H., Nguyen, H.V., Nguyen, H.L., Sauve, E., Nguyen, N.T.H., Chopo-Pizarro, A., Trimarsanto, H., Monsieurs, P., Guetens, P., and Nguyen, X.X. (2022). Novel highly-multiplexed AmpliSeq targeted assay for Plasmodium vivax genetic surveillance use cases at multiple geographical scales. Frontiers in Cellular and Infection Microbiology, 1154.

Koepfli, C., Rodrigues, P.T., Antao, T., Orjuela-Sánchez, P., Van Den Eede, P., Gamboa, D., Van Hong, N., Bendezu, J., Erhart, A., and Barnadas, C. (2015). Plasmodium vivax diversity and population structure across four continents. PLoS neglected tropical diseases 9, e0003872.

Lee, S., and Bahlo, M. (2016). moimix: an R package for assessing clonality in high-throughput sequencing data. moimix: an R package for assessing clonality in high-throughput sequencing data.

Lin, J.T., Patel, J.C., Kharabora, O., Sattabongkot, J., Muth, S., Ubalee, R., Schuster, A.L., Rogers, W.O., Wongsrichanalai, C., and Juliano, J.J. (2013). Plasmodium vivax isolates from Cambodia and Thailand show high genetic complexity and distinct patterns of P. vivax multidrug resistance gene 1 (pvmdr1) polymorphisms. The American journal of tropical medicine and hygiene 88, 1116–1123.

Manni, M., Berkeley, M.R., Seppey, M., Simão, F.A., and Zdobnov, E.M. (2021). BUSCO update: novel and streamlined workflows along with broader and deeper phylogenetic coverage for scoring of eukaryotic, prokaryotic, and viral genomes. Molecular Biology and Evolution 38, 4647–4654.

Minassian, A.M., Themistocleous, Y., Silk, S.E., Barrett, J.R., Kemp, A., Quinkert, D., Nielsen, C.M., Edwards, N.J., Rawlinson, T.A., and Lopez, F.R. (2021). Controlled human malaria infection with a clone of Plasmodium vivax with high quality genome assembly. JCI insight.

Mok, B.W., Ribacke, U., Sherwood, E., and Wahlgren, M. (2008). A highly conserved segmental duplication in the subtelomeres of Plasmodium falciparum chromosomes varies in copy number. Malaria journal 7, 1–9.

Neafsey, D.E., Galinsky, K., Jiang, R.H., Young, L., Sykes, S.M., Saif, S., Gujja, S., Goldberg, J.M., Young, S., and Zeng, Q. (2012). The malaria parasite Plasmodium vivax exhibits greater genetic diversity than Plasmodium falciparum. Nature genetics 44, 1046–1050.

Noviyanti, R., Miotto, O., Barry, A., Marfurt, J., Siegel, S., Thuy-Nhien, N., Quang, H.H., Anggraeni, N.D., Laihad, F., and Liu, Y. (2020). Implementing parasite genotyping into national surveillance frameworks: feedback from control programmes and researchers in the Asia–Pacific region. Malaria Journal 19.

Ntumngia, F.B., Thomson-Luque, R., De Menezes Torres, L., Gunalan, K., Carvalho, L.H., and Adams, J.H. (2016). A novel erythrocyte binding protein of Plasmodium vivax suggests an alternate invasion pathway into Duffy-positive reticulocytes. MBio 7, e01261-01216.

Nurk, S., Walenz, B.P., Rhie, A., Vollger, M.R., Logsdon, G.A., Grothe, R., Miga, K.H., Eichler, E.E., Phillippy, A.M., and Koren, S. (2020). HiCanu: accurate assembly of segmental duplications, satellites, and allelic variants from high-fidelity long reads. Genome research 30, 1291–1305.

Otto, T.D., Böhme, U., Sanders, M., Reid, A., Bruske, E.I., Duffy, C.W., Bull, P.C., Pearson, R.D., Abdi, A., and Dimonte, S. (2018). Long read assemblies of geographically dispersed Plasmodium falciparum isolates reveal highly structured subtelomeres. Wellcome Open Research 3.

Pacbio (2020a). “Application Note - Considerations for Using the Low and Ultra-Low DNA Input Workflows for Whole Genome Sequencing V2”.).

Pacbio (2020b). Ultra-Low DNA Input Workflow for SMRT Sequencing [Online]. Available: https://www.pacb.com/plant-animal-biology/introducing-the-ultra-low-input-protocol-for-smrt-sequencing/ [Accessed].

Pearson, R.D., Amato, R., Auburn, S., Miotto, O., Almagro-Garcia, J., Amaratunga, C., Suon, S., Mao, S., Noviyanti, R., and Trimarsanto, H. (2016). Genomic analysis of local variation and recent evolution in Plasmodium vivax. Nature genetics 48, 959.

Ponzi, M., Pace, T., Dore, E., and Frontali, C. (1985). Identification of a telomeric DNA sequence in Plasmodium berghei. The EMBO journal 4, 2991–2995.

Popovici, J., Roesch, C., Carias, L.L., Khim, N., Kim, S., Vantaux, A., Mueller, I., Chitnis, C.E., King, C.L., and Witkowski, B. (2020). Amplification of Duffy binding protein-encoding gene allows Plasmodium vivax to evade host anti-DBP humoral immunity. Nature communications 11, 1–8.

R Core Team (2019). “R: A language and environment for statistical computing”. (Vienna, Austria: R Foundation for Statistical Computing).

Rieckmann, K., Davis, D., and Hutton, D. (1989). Plasmodium vivax resistance to chloroquine? The Lancet 334, 1183–1184.

Sereika, M., Kirkegaard, R.H., Karst, S.M., Michaelsen, T.Y., Sørensen, E.A., Wollenberg, R.D., and Albertsen, M. (2022). Oxford Nanopore R10.4 long-read sequencing enables the generation of near-finished bacterial genomes from pure cultures and metagenomes without short-read or reference polishing. Nature Methods 19, 823–826.

Silva, S.R., Almeida, A.C.G., Da Silva, G.a.V., Ramasawmy, R., Lopes, S.C.P., Siqueira, A.M., Costa, G.L., Sousa, T.N., Vieira, J.L.F., and Lacerda, M.V.G. (2018). Chloroquine resistance is associated to multi-copy pvcrt-o gene in Plasmodium vivax malaria in the Brazilian Amazon. Malaria journal 17, 1–8.

Sriprawat, K., Kaewpongsri, S., Suwanarusk, R., Leimanis, M.L., Phyo, A.P., Snounou, G., Russell, B., Renia, L., and Nosten, F. (2009). Effective and cheap removal of leukocytes and platelets from Plasmodium vivax infected blood. Malaria journal 8, 1–7.

Steinbiss, S., Silva-Franco, F., Brunk, B., Foth, B., Hertz-Fowler, C., Berriman, M., and Otto, T.D. (2016). Companion: a web server for annotation and analysis of parasite genomes. Nucleic acids research 44, W29–W34.

Suwanarusk, R., Chavchich, M., Russell, B., Jaidee, A., Chalfein, F., Barends, M., Prasetyorini, B., Kenangalem, E., Piera, K., and Lek-Uthai, U. (2008). Amplification of pvmdr1 associated with multidrug-resistant Plasmodium vivax. The Journal of infectious diseases 198, 1558–1564.

Vernick, K.D., and Mccutchan, T.F. (1988). Sequence and structure of a Plasmodium falciparum telomere. Molecular and biochemical parasitology 28, 85–94.

Villena, F.E., Sanchez, J.F., Nolasco, O., Braga, G., Ricopa, L., Barazorda, K., Salas, C.J., Lucas, C., Lizewski, S.E., and Joya, C.A. (2022). Drug resistance and population structure of Plasmodium falciparum and Plasmodium vivax in the Peruvian Amazon. Scientific Reports 12, 1–11.

Wenger, A.M., Peluso, P., Rowell, W.J., Chang, P.-C., Hall, R.J., Concepcion, G.T., Ebler, J., Fungtammasan, A., Kolesnikov, A., Olson, N.D., Töpfer, A., Alonge, M., Mahmoud, M., Qian, Y., Chin, C.-S., Phillippy, A.M., Schatz, M.C., Myers, G., Depristo, M.A., Ruan, J., Marschall, T., Sedlazeck, F.J., Zook, J.M., Li, H., Koren, S., Carroll, A., Rank, D.R., and Hunkapiller, M.W. (2019). Accurate circular consensus long-read sequencing improves variant detection and assembly of a human genome. Nature Biotechnology 37, 1155–1162.

Who (2021). “World malaria report”. (Geneva: WHO).

Wood, D.E., Lu, J., and Langmead, B. (2019). Improved metagenomic analysis with Kraken 2. Genome biology 20, 1–13.

World Medical Association (2013). World Medical Association Declaration of Helsinki: ethical principles for medical research involving human subjects. Jama 310, 2191–2194.

Zeeshan, M., Tyagi, R.K., Tyagi, K., Alam, M.S., and Sharma, Y.D. (2014). Host-parasite interaction: selective Pv-fam-a family proteins of Plasmodium vivax bind to a restricted number of human erythrocyte receptors. The Journal of infectious diseases 211, 1111–1120.

