## Supplementary data for "A new *Plasmodium vivax* reference genome for South American isolates"

### Supplementary Figures

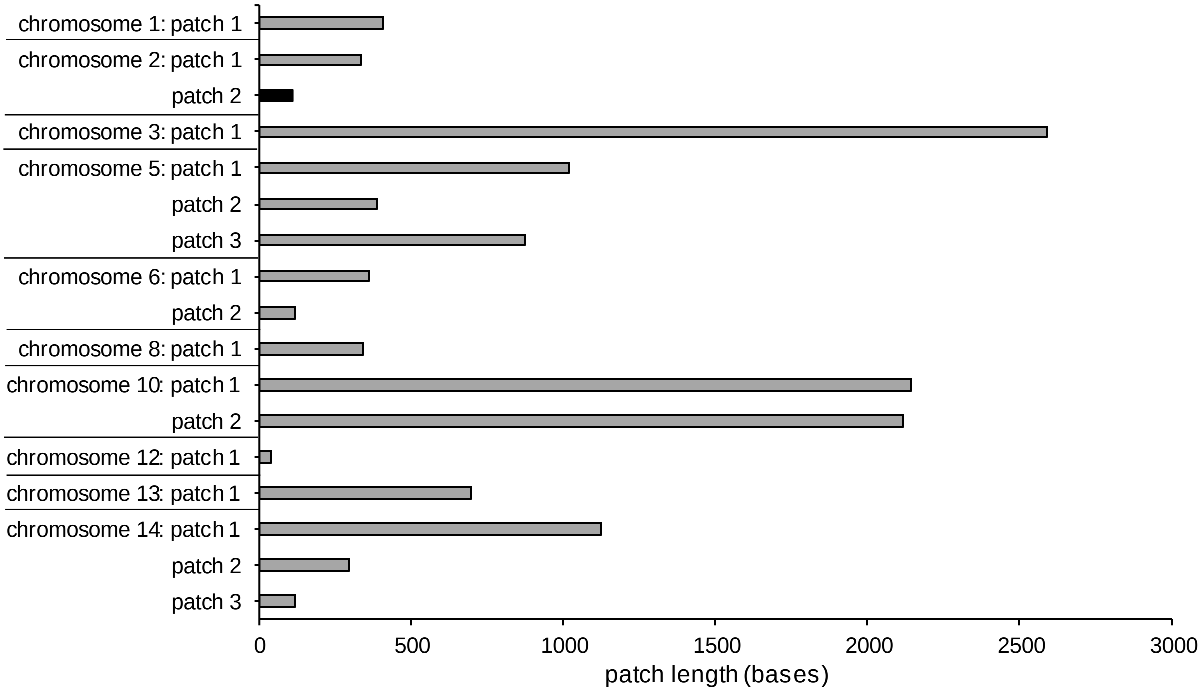


**Supplementary Figure 1.** Lengths of the patches that fill the gaps between the contigs scaffolded to the chromosomes. Grey bars indicate patches that are based on the PvW1 genome, black bars patches based on the PvP01 genome (when region was absent in PvW1). Chromosomes 4, 7, 9, 11 and the mitochondrial and apicoplast chromosome were fully assembled and contain no patched regions.

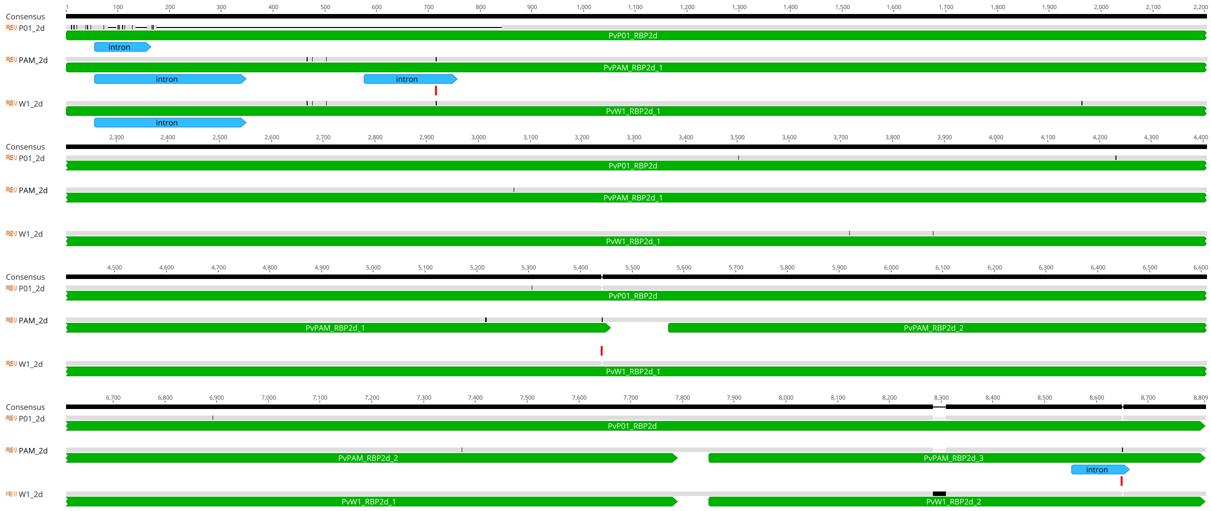


**Supplementary Figure 2.** Alignment of the *RBP2d* genes of PvP01 (1 pseudogene), PvPAM (3 coding genes) and PvW1 (2 coding genes). Indicated in red are 1 nt SNPs/indels in PvPAM which cause the difference in predicted introns and translation start/stop sites in comparison to PvW1.

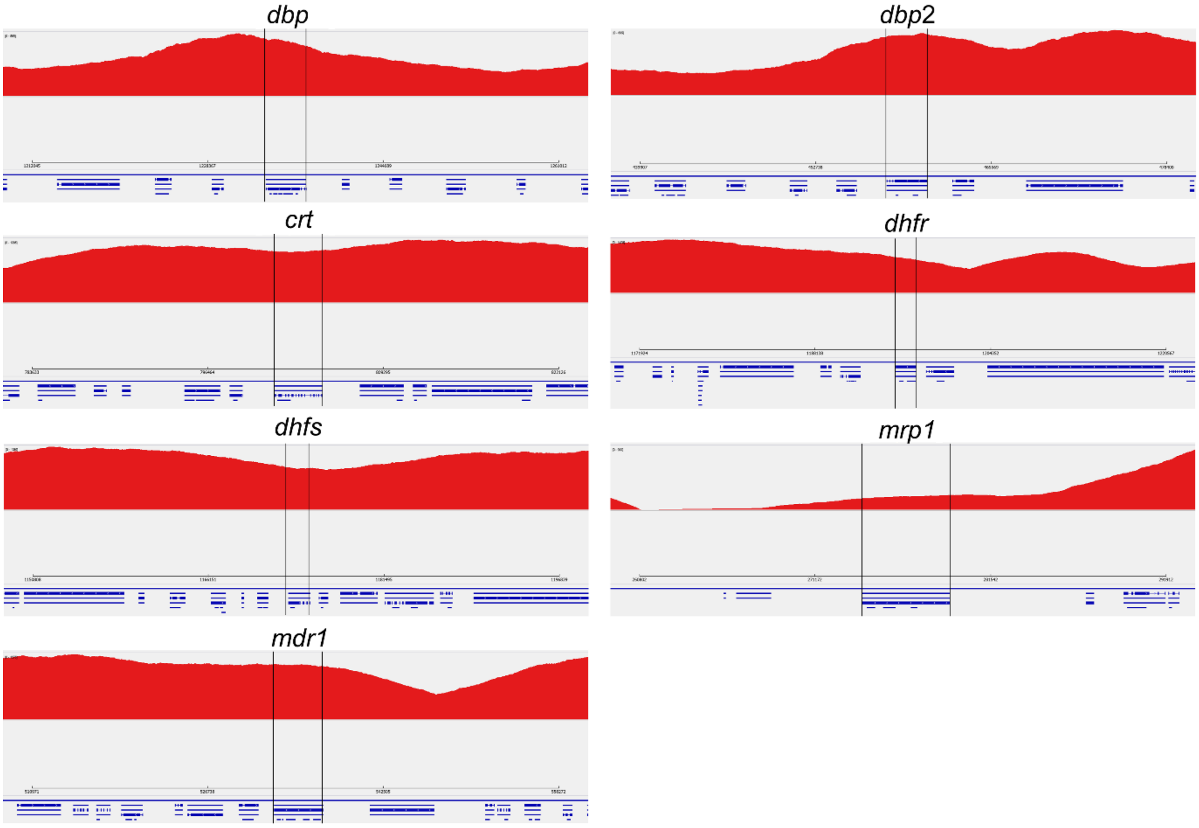


**Supplementary Figure 3.** Sashimi coverage plots created in IGV, showing PacBio reads of the PvPAM isolate mapped to the PvPAM reference genome. The gene under investigation for copy number variation is delineated with vertical black lines. Relative to the coverage level of the surrounding regions, no clear increase in coverage could be observed for any of the shown genes, which would have been indicative of copy number variation.

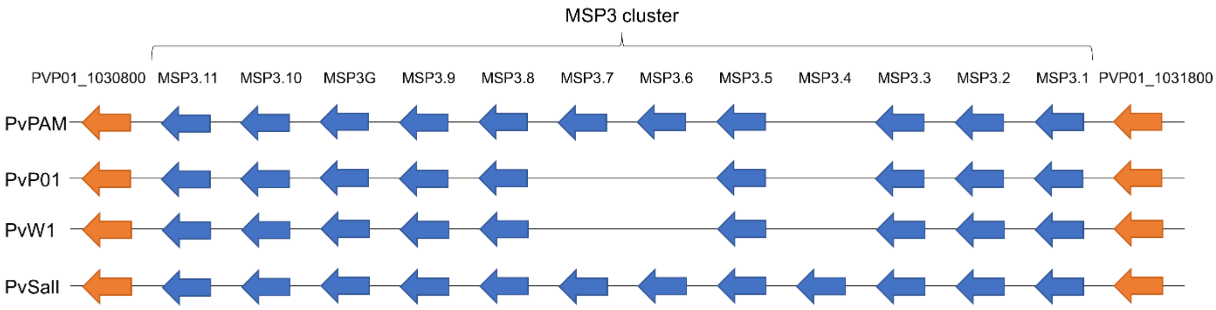


**Supplementary Figure 4.** MSP3 cluster organization in the PvPAM, PvP01, PvW1 and PvSalI reference genomes. Flanking genes (orange) of the MSP3 cluster are present in all isolates.

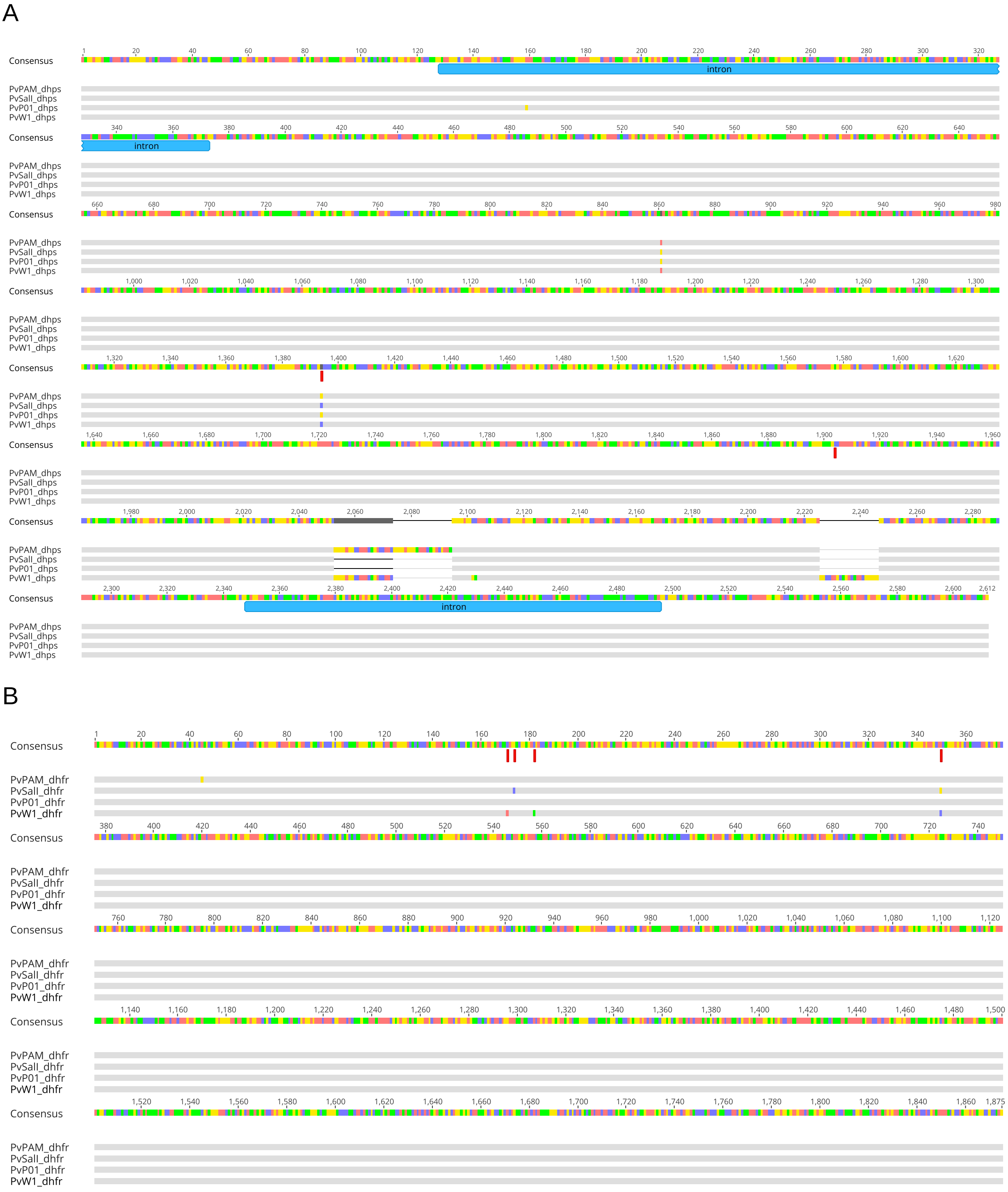
A
Consensus
intron
PvPAM_dhps
PvSall_dhps
PvP01_dhps
PvW1_dhps
B
PvPAM_dhfr
PvSall_dhfr
PvP01_dhfr
PvW1_dhfr

C

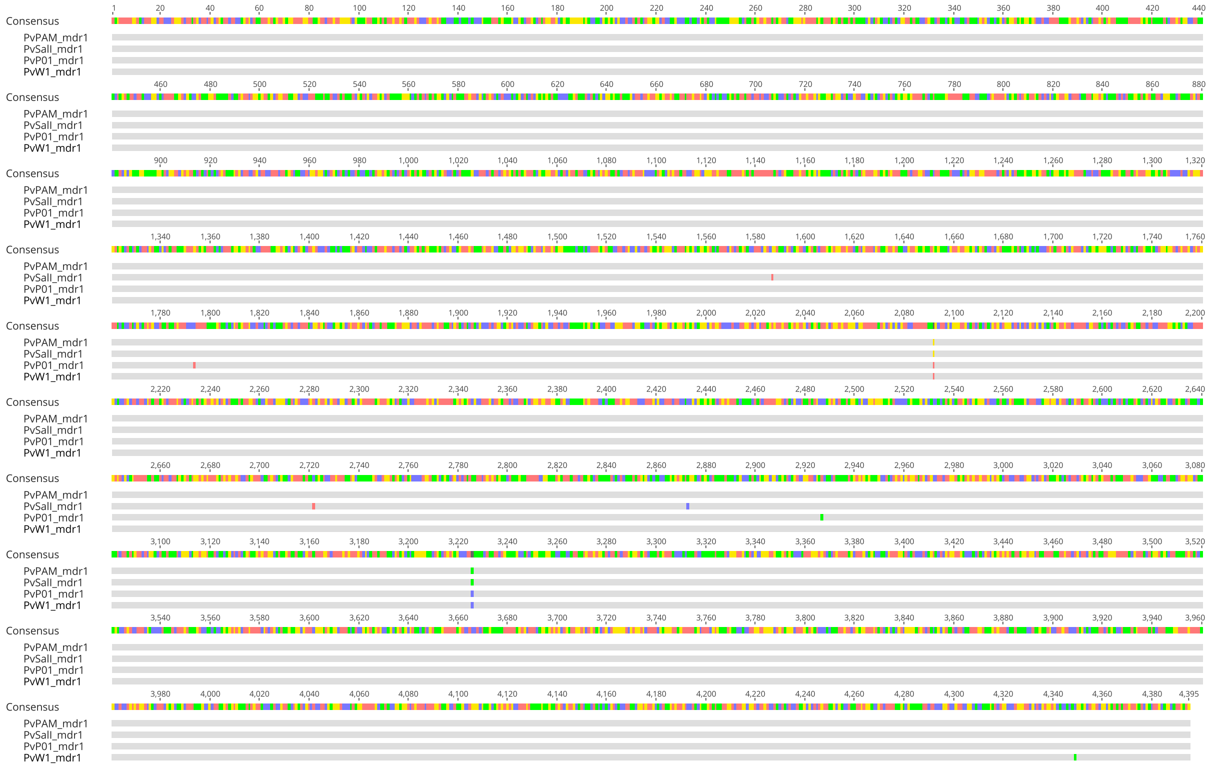


D

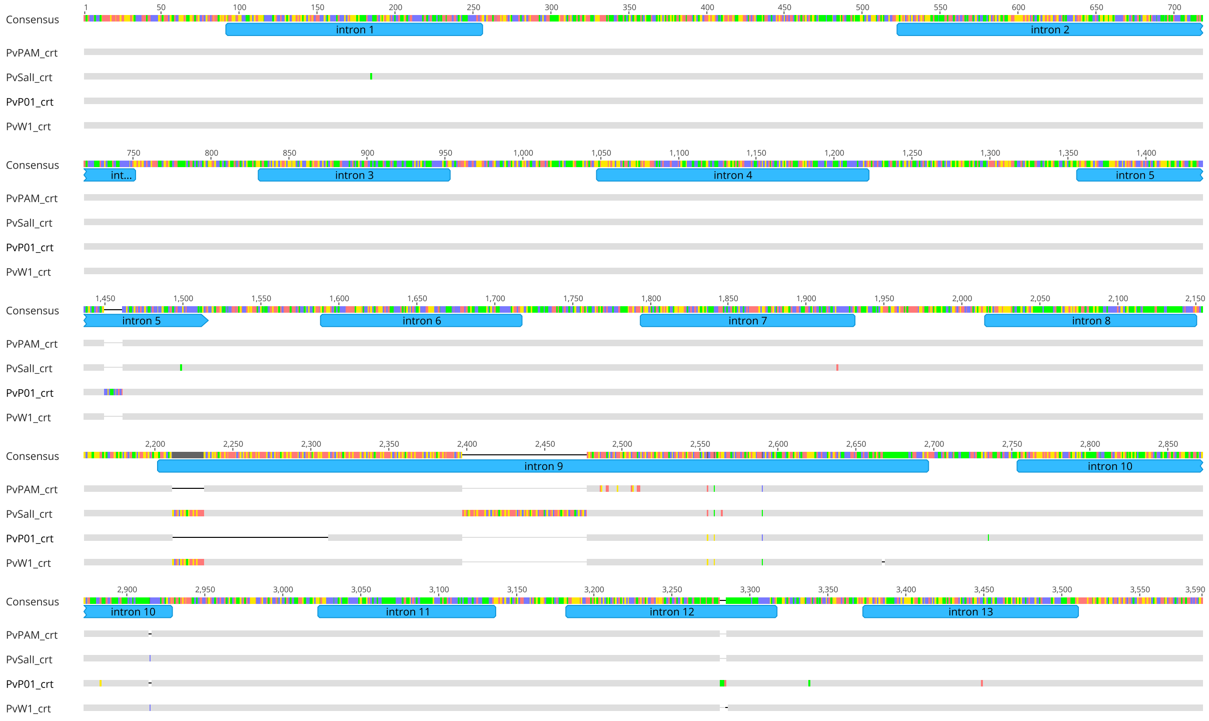


**Supplementary Figure 5.** Multiple sequence DNA alignments of the PvPAM, PvSalI, PvP01 and PvW1 *dhps* (**A**), *dhfr* (**B**), *mdr1* (**C**) and *crt* (**D**) genes, each shown in the sense direction (5’ to 3’). Resistance-associated SNPs are annotated as red blocks below the consensus sequence, introns are shown as blue blocks. Sequence differences are highlighted in colour, conserved bases are shown in grey. Deletions are shown as horizontal black lines. Alignments were made with MUSCLE.

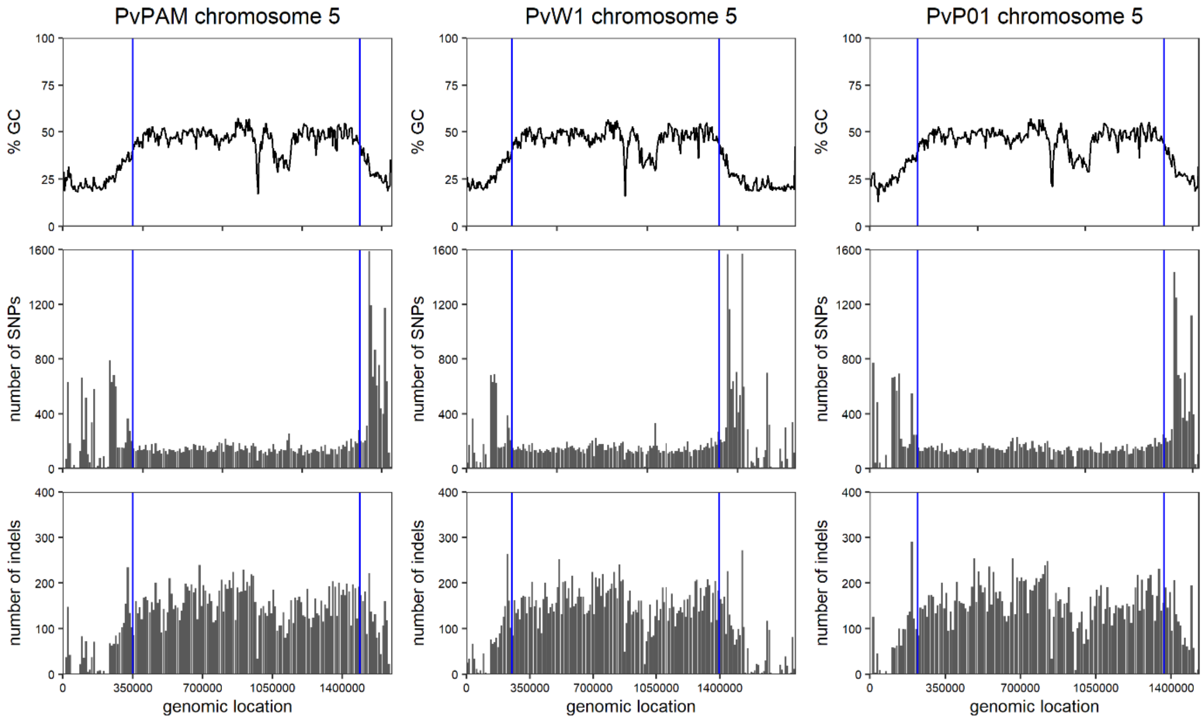


**Supplementary Figure 6.** Number of SNPs (middle row) and indels (bottom row) when Illumina reads from South American samples (N=354) are mapped to chromosome 5 of PvPAM, PvW1 (contig CAJZCX010000010) and PvP01. The top row plots show the GC content of chromosome 5, and blue vertical lines mark the border between the core and subtelomeric genome as determined in this study.

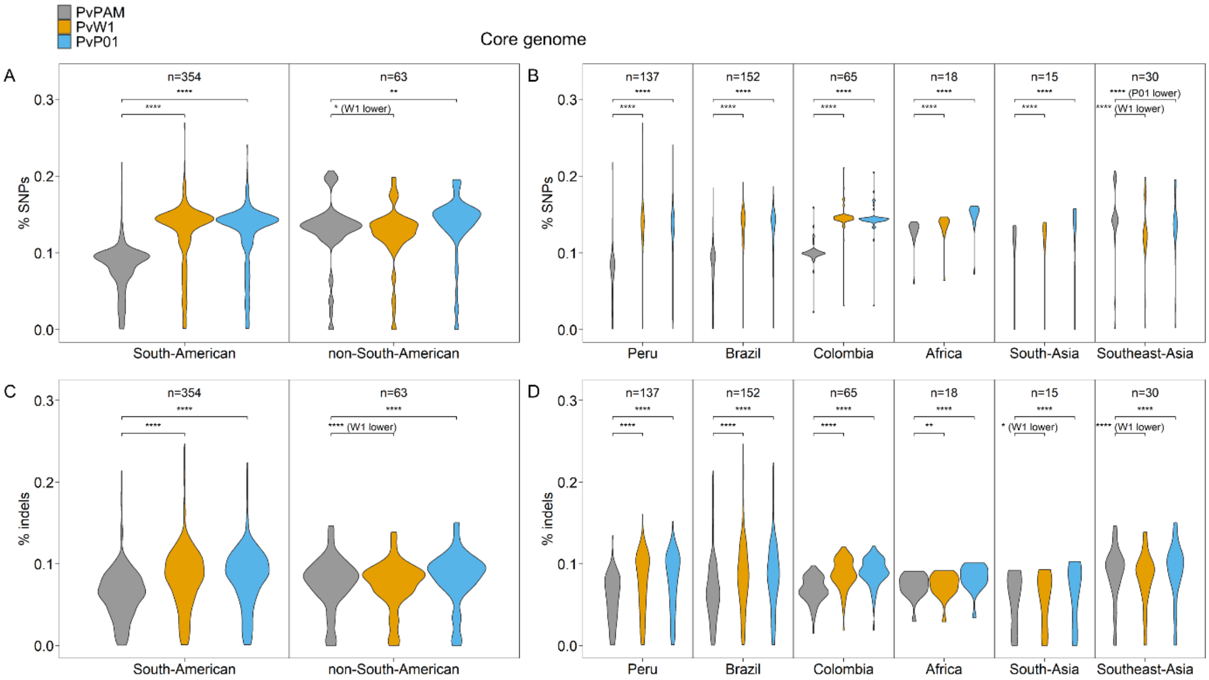


**Supplementary Figure 7.** Violin plots showing the percentage of variants (variants/bp) in the core genome for Illumina WGS reads from different geographical regions mapped to the PvPAM, PvW1 or PvP01 reference genome. **A.** % of SNPs in South American vs non-South American samples; **B.** % of SNPs per geographical region; **C.** % of indels in South American vs non-South American samples; **D.** % of indels per geographical region. * $p<0.05$, ** $p<0.01$, *** $p<0.001$, **** $p<0.0001$ (paired Wilcoxon signed-rank test). If significant, mapping to PvPAM results in a lower % of variants, unless stated otherwise alongside the significance value. It is not possible to compare the % of variants between the South American and non-South American region, since those reads originate from different samples. African samples originate from Eritrea (5), Ethiopia (5), Sudan (5) and Uganda (3), South Asian samples from Afghanistan (5), India (5) and Pakistan (5), and Southeast Asian samples from Cambodia (5), Malaysia (5), Myanmar (5), Papua New Guinea (5), Thailand (5) and Vietnam (5). n=number of Illumina WGS samples.

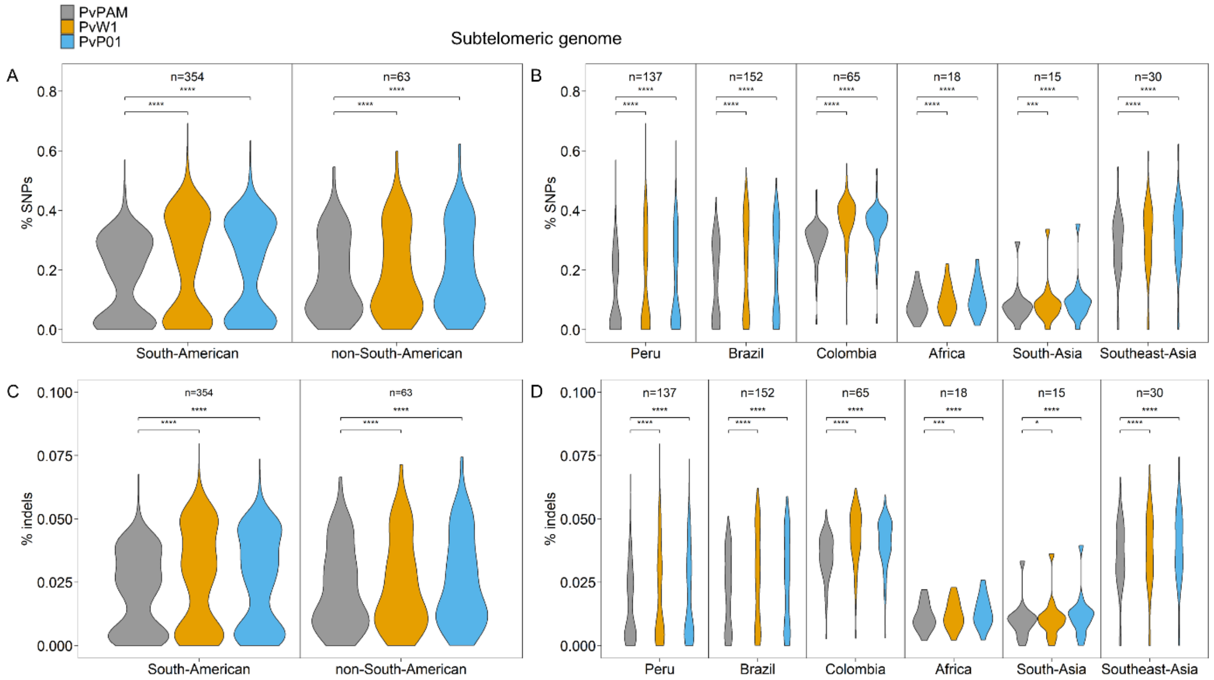


**Supplementary Figure 8.** Violin plots showing the percentage of variants (variants/bp) in the subtelomeric genome for Illumina WGS reads from different geographical regions mapped to the PvPAM, PvW1 or PvP01 reference genome. **A.** % of SNPs in South American vs non-South American samples; **B.** % of SNPs per geographical region; **C.** % of indels in South American vs non-South American samples; **D.** % of indels per geographical region. * $p<0.05$, ** $p<0.01$, *** $p<0.001$, **** $p<0.0001$ (paired Wilcoxon signed-rank test). If significant, mapping to PvPAM results in a lower % of variants, unless stated otherwise alongside the significance value. It is not possible to compare the % of variants between the South American and non-South American region, since those reads originate from different samples. African samples originate from Eritrea (5), Ethiopia (5), Sudan (5) and Uganda (3), South Asian samples from Afghanistan (5), India (5) and Pakistan (5), and Southeast Asian samples from Cambodia (5), Malaysia (5), Myanmar (5), Papua New Guinea (5), Thailand (5) and Vietnam (5). n=number of Illumina WGS samples.

### Supplementary Tables

**Supplementary Table 1.** Predicted function and Pfam domain ID’s of newly identified genes (excluding pseudogenes) in the PvPAM assembly, that are not member of an orthology group containing a PvP01 (pseudo)gene.

| Number of genes | Description | Pfam ID |
|---|---|---|
| 163 | hypothetical protein | NA |
| 60 | hypothetical protein, conserved | NA |
| 238 | Plasmodium vivax Vir protein | PF05795 |
| 17 | Protein of unknown function | PF12420 |
| 5 | Plasmodium RESA N-terminal | PF09687 |
| 1 | PH domain/Oxysterol-binding protein | PF00169, PF01237 |
| 1 | Plasmodium variant antigen protein Cir/Yir/Bi | PF06022 |
| 1 | Tumour-associated protein/Protein of unknown function | PF09746, PF12420 |
| 1 | ubiE/COQ5 methyltransferase family/Tellurite resistance protein TehB/Methyltransferase small domain/Methyltransferase domain containing protein | PF01209, PF03848, PF05175, PF13489, PF13847, PF13649, PF08242, PF08241 |

**Supplementary Table 2.** Overview of the PvRBP genes in PvPAM, PvP01, PvSalI and PvW1. Partial (shorter) genes are in italic, pseudogenes have a grey background.

| Gene name | PvPAM | PvP01 (PlasmoDB, v51) | PvSalI (PlasmoDB, v45) | PvW1 (PlasmoDB, v60) |
|---|---|---|---|---|
| RBP1a | PVPAM_070008100 | PVP01_0701200 | PVX_098585 | PVW1_070007700 |
| RBP1b | PVPAM_070008000 | PVP01_0701100 | PVX_098582, PVX_125738 | PVW1_070007600 |
| RBP2a | PVPAM_140009900 | PVP01_1402400 | PVX_121920 | PVW1_140008600 |
| RBP2b | PVPAM_080012700 | PVP01_0800700 | PVX_094255 | PVW1_080006000 |
| RBP2c | PVPAM_050042100 | PVP01_0534300 | PVX_090325 | PVW1_050039600 |
| RBP1-p1 | / | *PVP01_0010770* | / | / |
| RBP2-p1 | *PVPAM_050042200* | *PVP01_0534400* | *PVX_090330* | *PVW1_050039800* |
| RBP2-p2 | / | / | *PVX_101590* | *On chromosome 14, not annotated* |
| RBP2d | *PVPAM_140080500, PVPAM_140080400, PVPAM_140080300 ** | PVP01_1471400 | PVX_101585 | *PVW1_140078000, PVW1_140077900 ** |
| RBP2e | PVPAM_070007400 | PVP01_0700500 | / | *PVW1_070006900 ** |
| RBP3 | PVPAM_140078200 | PVP01_1469400 | PVX_101495 | On chromosome 14, not annotated |

* Caution is needed with predicted gene annotations: depending on the location of translation start/stop sites and introns, they could also be a pseudogene (in case of RBP2d, the combination of the different partial genes could be 1 pseudogene).

**Supplementary Table 3.** Overview of the antifolate resistance-associated *dhps* and *dhfr* mutations ('x') in PvPAM, PvSalI, PvP01 or PvW1.

| | | PvPAM | PvSalI | PvP01 | PvW1 | References |
|---|---|---|---|---|---|---|
| DHPS | A383G | x | | x | | (Korsinczky et al., 2004;Imwong et al., 2008) |
| | A553G | | | | | |
| DHFR | F57L | | | | x | (Tjitra et al., 2002;Hastings et al., 2005;Auliff et al., 2006;Marfurt et al., 2008;Rungsihirunrat et al., 2008;Zakeri et al., 2009) |
| | S58R | x | | x | x | |
| | T61M | | | | x | |
| | S117T | | | | x | |
| | S117N | x | | x | | |

**Supplementary Table 4.** BUSCO results for PvPAM, PvP01 (PlasmoDB, v51) and PvW1 (PlasmoDB, v60).

| | **PvPAM** | **PvP01** | **PvW1** |
|---|---|---|---|
| Total BUSCO groups searched | 3642 | | |
| Complete BUSCOs | 3526 | 3530 | 3525 |
| - Complete and single-copy BUSCOs | 3526 | 3530 | 3525 |
| - Complete and duplicated BUSCOs | 0 | 0 | 0 |
| Fragmented BUSCOs | 30 | 25 | 31 |
| Missing BUSCOs | 86 | 87 | 86 |

### References


Auliff, A., Wilson, D.W., Russell, B., Gao, Q., Chen, N., Le Ngoc, A., Maguire, J., Bell, D., O'neil, M.T., and Cheng, Q. (2006). Amino acid mutations in Plasmodium vivax DHFR and DHPS from several geographical regions and susceptibility to antifolate drugs. *The American journal of tropical medicine and hygiene* 75**,** 617-621.

Hastings, M.D., Maguire, J.D., Bangs, M.J., Zimmerman, P.A., Reeder, J.C., Baird, J.K., and Sibley, C.H. (2005). Novel Plasmodium vivax dhfr alleles from the Indonesian Archipelago and Papua New Guinea: association with pyrimethamine resistance determined by a Saccharomyces cerevisiae expression system. *Antimicrobial agents and chemotherapy* 49**,** 733-740.

Imwong, M., Pukrittayakamee, S., Pongtavornpinyo, W., Nakeesathit, S., Nair, S., Newton, P., Nosten, F., Anderson, T.J., Dondorp, A., and Day, N.P. (2008). Gene amplification of the multidrug resistance 1 gene of Plasmodium vivax isolates from Thailand, Laos, and Myanmar. *Antimicrobial agents and chemotherapy* 52**,** 2657-2659.

Korsinczky, M., Fischer, K., Chen, N., Baker, J., Rieckmann, K., and Cheng, Q. (2004). Sulfadoxine resistance in Plasmodium vivax is associated with a specific amino acid in dihydropteroate synthase at the putative sulfadoxine-binding site. *Antimicrobial agents and chemotherapy* 48**,** 2214-2222.

Marfurt, J., De Monbrison, F., Brega, S., Barbollat, L., Müller, I., Sie, A., Goroti, M., Reeder, J.C., Beck, H.-P., and Picot, S. (2008). Molecular markers of in vivo Plasmodium vivax resistance to amodiaquine plus sulfadoxine-pyrimethamine: mutations in pvdhfr and pvmdr1. *The Journal of Infectious Diseases* 198**,** 409-417.

Rungsihirunrat, K., Sibley, C.H., Mungthin, M., and Na-Bangchang, K. (2008). Geographical distribution of amino acid mutations in Plasmodium vivax DHFR and DHPS from malaria endemic areas of Thailand. *The American journal of tropical medicine and hygiene* 78**,** 462-467.

Tjitra, E., Baker, J., Suprianto, S., Cheng, Q., and Anstey, N.M. (2002). Therapeutic efficacies of artesunate-sulfadoxine-pyrimethamine and chloroquine-sulfadoxine-pyrimethamine in vivax malaria pilot studies: relationship to Plasmodium vivax dhfr mutations. *Antimicrobial agents and chemotherapy* 46**,** 3947-3953.

Zakeri, S., Motmaen, S.R., Afsharpad, M., and Djadid, N.D. (2009). Molecular characterization of antifolates resistance-associated genes,(dhfr and dhps) in Plasmodium vivax isolates from the Middle East. *Malaria Journal* 8**,** 1-9.
